# Myosin forces elicit an F-actin structural landscape that mediates mechanosensitive protein recognition

**DOI:** 10.1101/2024.08.15.608188

**Authors:** Ayala G. Carl, Matthew J. Reynolds, Pinar S. Gurel, Donovan Y.Z. Phua, Xiaoyu Sun, Lin Mei, Keith Hamilton, Yasuharu Takagi, Alex J. Noble, James R. Sellers, Gregory M. Alushin

## Abstract

Cells mechanically interface with their surroundings through cytoskeleton-linked adhesions, allowing them to sense physical cues that instruct development and drive diseases such as cancer. Contractile forces generated by myosin motor proteins mediate these mechanical signal transduction processes through unclear protein structural mechanisms. Here, we show that myosin forces elicit structural changes in actin filaments (F-actin) that modulate binding by the mechanosensitive adhesion protein α-catenin. Using correlative cryo-fluorescence microscopy and cryo-electron tomography, we identify F-actin featuring domains of nanoscale oscillating curvature at cytoskeleton-adhesion interfaces enriched in zyxin, a marker of actin-myosin generated traction forces. We next introduce a reconstitution system for visualizing F-actin in the presence of myosin forces with cryo-electron microscopy, which reveals morphologically similar superhelical F-actin spirals. In simulations, transient forces mimicking tugging and release of filaments by motors produce spirals, supporting a mechanistic link to myosin’s ATPase mechanochemical cycle. Three-dimensional reconstruction of spirals uncovers extensive asymmetric remodeling of F-actin’s helical lattice. This is recognized by α-catenin, which cooperatively binds along individual strands, preferentially engaging interfaces featuring extended inter-subunit distances while simultaneously suppressing rotational deviations to regularize the lattice. Collectively, we find that myosin forces can deform F-actin, generating a conformational landscape that is detected and reciprocally modulated by a mechanosensitive protein, providing a direct structural glimpse at active force transduction through the cytoskeleton.

## Main

Myosin motor proteins hydrolyze ATP to apply contractile forces to cytoskeletal actin filaments (F-actin)^1,2^ linked to plasma-membrane spanning adhesion complexes^3,4^, enabling cells to mechanically interface with their local tissue microenvironments. Actin-myosin contractility is critical for cell migration^5,6^, cell division^7^, and multi-cellular tissue dynamics^8^, fundamental processes required for proper morphogenesis and tissue homeostasis that are frequently dysregulated in developmental diseases and cancer^9–11^. In addition to directly shaping cells and tissues, contractile forces are prominently associated with the transduction of mechanical cues into biochemical signaling pathways that govern cell behavior^12–14^. How piconewton-scale forces generated by myosins initiate downstream mechanical signal transduction remains broadly unclear at the protein structural level.

Beyond serving as a general force-generating apparatus for mechanical signaling pathways, the cytoskeleton can itself mediate force transduction through mechanically-regulated binding interactions between F-actin and actin-binding proteins (ABPs)^15–18^. The intrinsic actin-binding activity of canonical ABPs can be modulated by force, including regulators of F-actin assembly^19–25^ and disassembly^26,27^, crosslinkers^28–30^, and cell adhesion proteins^31–34^, with the critical cell-cell adhesion protein α-catenin specifically displaying enhanced binding to F-actin in the presence of active myosin force generation^34^. Collectively, these factors contribute to the force sensitive dynamics of cytoskeletal networks underlying cell migration^35,36^ and adhesion^37^. Additionally, several proteins containing LIN-11, Isl-1, and MEC-3 (LIM) domains solely bind F-actin in the presence of force^38,39^, thereby localizing to the cytoskeleton in the presence of active contractility^40–42^. This force-activated actin binding activity facilitates repair of mechanical damage to the cytoskeleton by zyxin^43^ and governs mechanosensitive nuclear localization of the gene expression control protein FHL2^14,38^. How these proteins detect the presence of force on F-actin is unknown.

F-actin features intrinsic structural polymorphism^44–46^, as well as specific conformational states when bound by ABPs such as the F-actin disassembly factor cofilin^47–49^, suggesting force could modulate F-actin structure to regulate ABP engagement^15^. Indirect reporters of actin subunit structural dynamics within the filament^45^, as well as X-ray diffraction studies of contracting muscle fibers^50,51^, have suggested F-actin structural rearrangements occur in the presence of myosin activity. Cryo-electron microscopy (cryo-EM) structural studies of F-actin–myosin complexes performed under saturation binding conditions have furthermore visualized subtle alterations in actin subunit conformation at specific stages of the motor’s mechanochemical cycle^52–55^. However, it is unclear whether and how these local changes at the actin-myosin interface propagate along the filament to modulate additional ABP binding sites. We recently reported that filament bending elicited by thermal fluctuations and fluid flow substantially alters F-actin structure, remodeling inter-subunit interfaces that are frequently engaged by ABPs^56^. Whether active myosin force generation also substantively remodels F-actin, and how ABPs discriminate mechanically- excited F-actin structural states through binding contacts, remain to be determined.

## Results

### Oscillatory F-actin domains are present in cell adhesions marked by zyxin

To investigate whether myosin contractility can modulate F-actin structure in cells, we pursued cryo-electron tomography (cryo-ET) studies. Zyxin’s subcellular localization pattern was recently reported to be sufficient to infer the pattern of traction forces exerted through cell-extracellular matrix (focal) adhesions^57^, consistent with the protein’s capacity to detect and bind tensed F-actin. We therefore employed zyxin as a marker of adhesion sites featuring high mechanical load. We examined Ptk2 cells stably expressing doxycycline inducible zyxin-mNeonGreen and the F-actin label fTractin-mScarlet. After extensive optimization (Methods; Fig. S1A), we found these cells to be thin enough to directly image adhesion-cytoskeleton interfaces without cryo-focused ion beam milling. To increase the contractility of cells cultured on holey carbon cryo-EM substrates, we treated them with Rho Activator II (Methods), a RhoA activating toxin.

Using correlative cryo-fluorescence microscopy, we targeted zyxin-enriched adhesions at the cell periphery, focusing on positions suitable for cryo-ET data collection (Fig. 1a; Fig. S1b,c; Video S1). Semantic segmentation of tomograms showed an abundance of actin filaments in both co-linear bundles (Fig. S1b), as previously reported in a cryo-ET study of focal adhesions marked by paxillin^58^, as well as more disorganized networks (Fig. 1b, left; Fig. S1c). Many filaments in both of these types of networks exhibited substantial curvature (Fig. 1b, left; Fig. S1b,c). Strikingly, this including individual filaments featuring domains of sharply oscillating curvature spanning ∼300-400 nanometers (corresponding to ∼100-150 subunits), which were continuous with canonical straight F-actin (Fig. 1b, right; Fig. S1b,c). As this morphology is distinct from the uniplanar curvature of bent filaments found at sites of propulsive actin polymerization against membranes^59–61^, we hypothesized it could be specifically evoked by myosin forces.

**Fig. 1:**
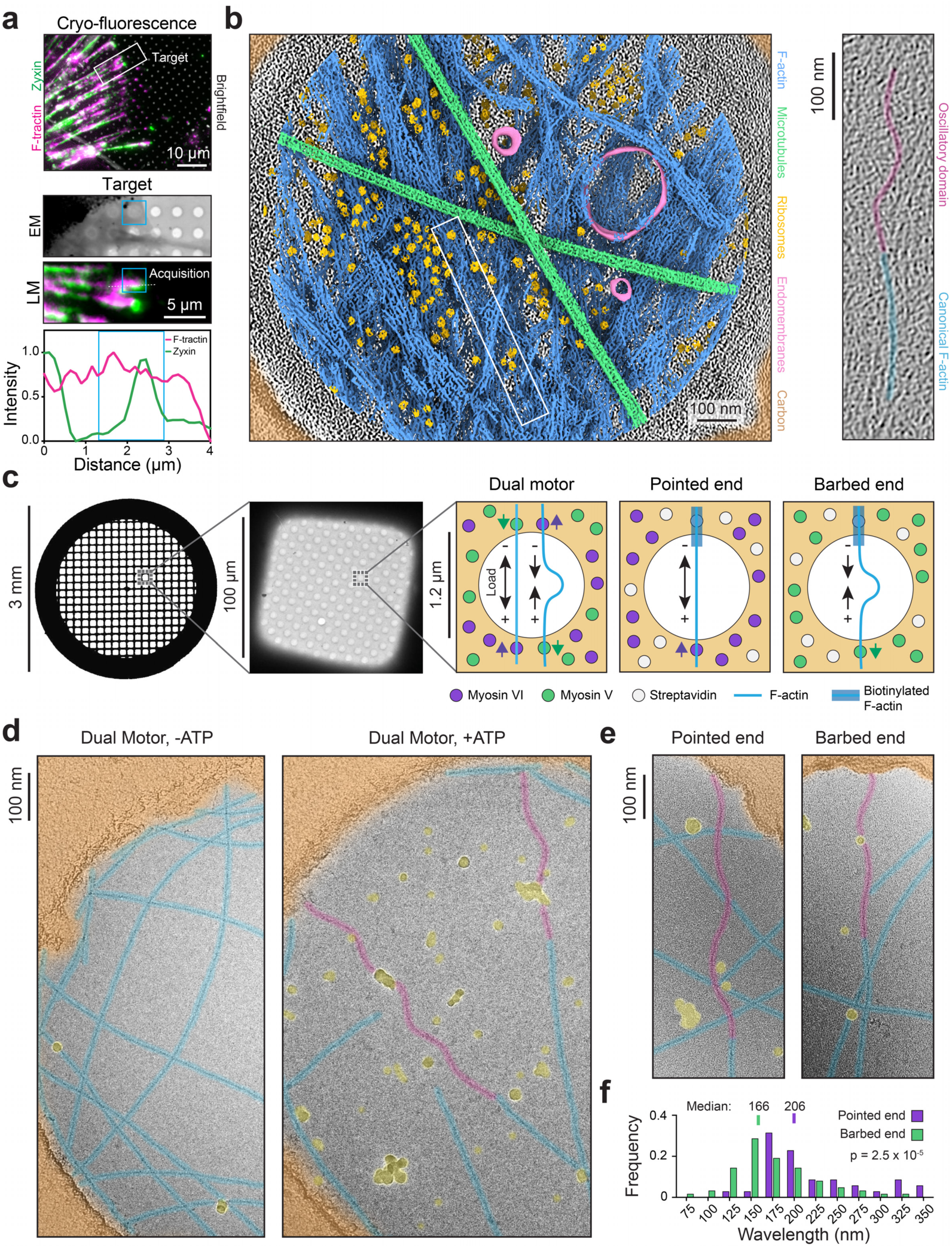
Myosin forces evoke oscillatory domains in F-actin. **a**, Top: Low-magnification cryo-light microscopy (LM) image of Ptk2 cell, highlighting targeted adhesion. Middle: Medium magnification correlation between cryo-electron microscopy (EM) and LM, highlighting site of tomogram acquisition. Bottom: Fluorescence intensity scan along dashed line. Blue box indicates acquisition area, which is enriched in zyxin. **b**, Left: segmented tomogram. Right: False colored 5.1 nm thick projection of boxed area, highlighting F-actin oscillatory domain. **c**, Schematic of myosin force reconstitution assay. **d**, False-colored cryo-EM images of dual motor reconstitution in the presence and absence of ATP. Oscillatory domains, magenta; canonical F-actin, blue; carbon film, orange; ice contamination, yellow. **e**, Cryo-EM images of oscillatory domains formed in single-motor conditions, false colored as in d. **f**, Quantification of oscillatory domain wavelengths in the pointed end directed (n = 35) and barbed end directed (n = 63) conditions, from N = 2 independent experiments. Distributions were compared with an unpaired two-tailed Mann-Whitney test.

### Myosin forces evoke a superhelical F-actin conformation

To examine whether myosin forces directly modulate F-actin conformation, we reconstituted myosin activity on holey carbon cryo-EM substrates (Methods), adapting an approach we have previously used to analyze force-activated F-actin binding proteins with fluorescence microscopy^34,38^. In this “dual motor” assay, a mixture of plus (“barbed”)-end directed myosin-5 and minus (“pointed”)-end directed myosin-6 are immobilized on a substrate, where they engage in a tug-of-war in the presence of ATP to apply mechanical stress to surface-adjacent F-actin (Fig. 1c). In the presence of myosin-5 or myosin-6 individually, fluorescence microscopy confirmed ATP-dependent gliding of F-actin across cryo-EM substrates (Video S2), while the dual motor condition results in desultory motions and breakage events (Video S3), consistent with filaments coming under mechanical load.

We next plunge-froze specimens to arrest motor dynamics and imaged them by cryo-EM (Methods). As the motors are anchored to the carbon film, imaging in holes facilitates visualizing force-dependent rearrangements which propagate along filaments, distal from local allosteric effects at motor binding sites. Consistent with our cellular cryo-ET data, we observed the appearance of oscillatory F-actin domains that spanned hundreds of nanometers in the presence of active force generation, continuous with canonical straight F-actin domains within the same filament (Fig. 1d). These domains were morphologically similar to those we observed in cells, with a median wavelength of 159 nm (Figure S2a). In addition to finding oscillatory domains in filaments spanning holes (Fig. S2b), a configuration in which they can bear load, we also observe oscillatory domains in non-load bearing configurations where they project into holes from the ends of filaments that contact the carbon film along a single edge (Figure S2c). We interpret these to represent remnants of mechanical severing events, as we observed with fluorescence microscopy (Video S3). This suggests oscillatory F-actin domains have the capacity to persist after filament breakages, when force has dissipated.

As the dual motor system is anticipated to produce a complex distribution of forces, we next sought to determine the role of motor directionality in oscillatory domain formation. We modified our system by polymerizing F-actin from biotinylated seeds (Methods), producing filaments with biotinylated regions at their pointed ends. In the presence of surface-anchored streptavidin, immobilized filaments were then exposed to either myosin-6 (“pointed-end directed force”) or myosin-5 (“barbed-end directed force”) individually. Fluorescence microscopy of the pointed-end directed force condition showed straight filaments which began gliding after mechanical ruptures, while the barbed-end directed force condition produced micron-scale buckling, consistent with the anticipated force distribution in each preparation (Video S4). Unexpectedly, cryo-EM images revealed the formation of oscillatory domains in both conditions (Fig. 1e). Oscillatory domains formed in the pointed-end directed force condition featured a significantly longer median wavelength (206 nm) than those formed in the barbed-end directed force condition (166 nm), but they were otherwise morphologically similar (Fig. 1f). These observations suggest that the formation of oscillatory domains represents an intrinsic response of F-actin to myosin forces, which is insensitive to motor directionality.

To gain more detailed insight into the ultrastructure of oscillatory domains, we pursued cryo-ET studies of both the barbed-end directed and pointed-end directed force conditions. Visual inspection suggested that oscillatory domains exhibited three-dimensional character (Fig. 2a; Video S5). However, degraded resolution in the Z-dimension due to incomplete tilt angle coverage (the “missing wedge” artifact) precluded detailed analysis. To overcome this limitation, we applied a neural-network–based tomogram denoising approach we recently developed^62^ (Methods), which minimized the missing wedge artifact and enabled direct filament tracing for quantification (Fig. 2b). Oscillatory regions approximately 300 nm in length were identified and subjected to principal component analysis (PCA) to decompose oscillations into orthogonal planes oriented along each filament’s long axis (Methods). Alignment of these filament trace projections revealed a consistent phase offset of approximately a quarter wavelength between each domain’s oscillatory components, despite substantial morphological variability (Fig. 2c,d). This observation is consistent with oscillatory domains adopting a superhelical spiral morphology which corkscrews around the filament axis.

**Fig. 2:**
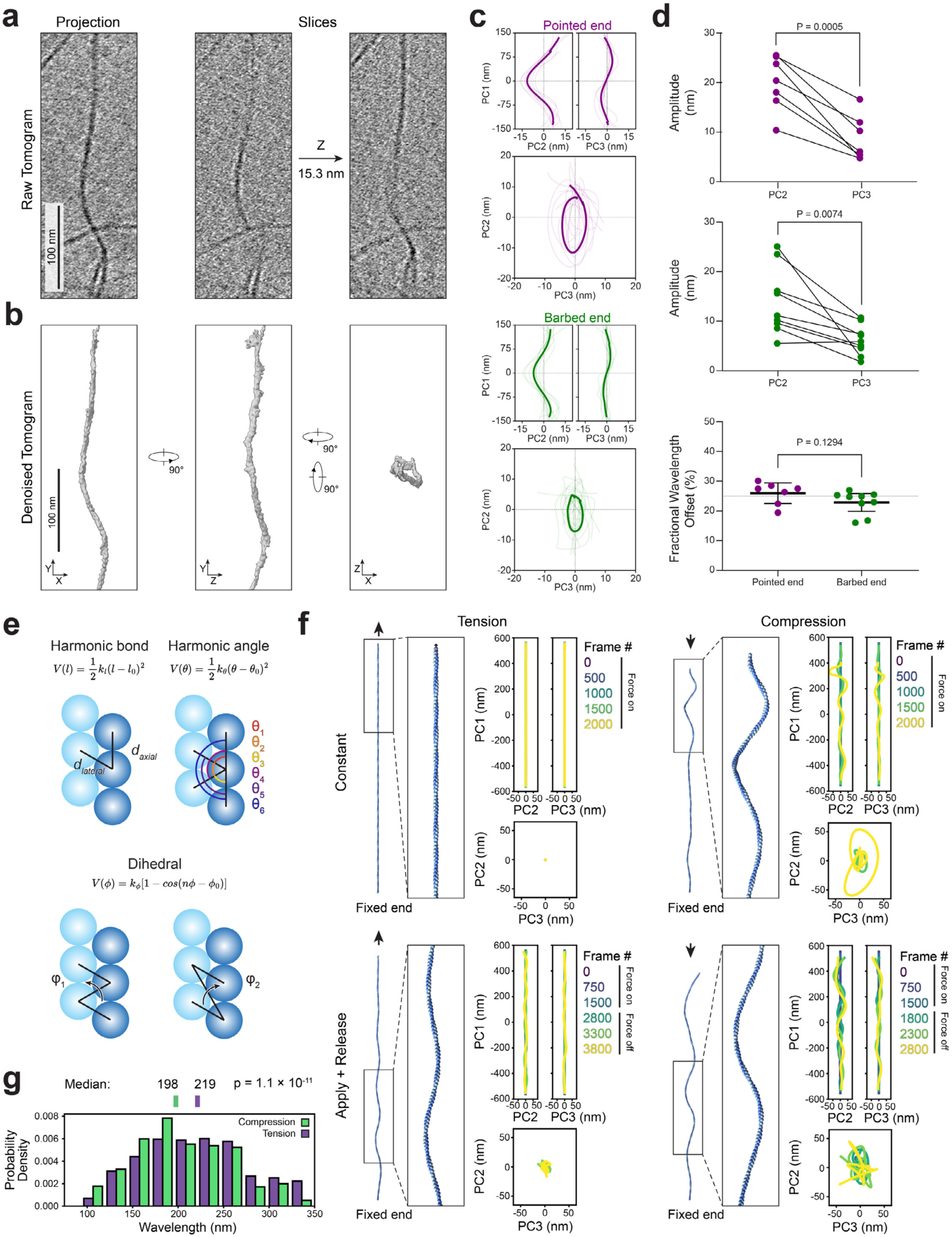
Transient force produces superhelical F-actin spirals. **a**, Left: projection and right: serial slices of an F-actin oscillatory domain tomogram (pointed end directed force condition). **b**, Orthogonal views of denoised density from tomogram in a, highlighting spiraling oscillations in both XY and XZ dimensions. **c**, Aligned projection views along principal components (PC) of filament traces with spiral character in pointed end directed (purple, top, n = 7) and barbed end directed (green, bottom, n = 9) force conditions. Transparent lines represent individual filament traces, while solid lines represent averages of aligned traces. **d**, Quantitation of data from c. Top and middle: Analysis of filament trace amplitudes in PC1-PC2 vs. PC1-PC3 planes from pointed-end (top) and barbed-end force conditions (middle), compared by paired t-test. Bottom: fractional wavelength offset between trace projections in PC1-PC2 vs. PC1-PC3 planes; force conditions were compared by unpaired t-test. **e**, Schematic of potentials used in coarse-grained molecular dynamics simulations. **f**, Representative simulation snapshots and principal component analyses of key frames in indicated conditions (constant force protocol, or apply then release force protocol, for both tension and compression). BE: barbed end; PE: pointed end. **g**, Quantification of oscillatory domain wavelengths from the apply then release force protocol. Conditions were compared by an unpaired t-test.

Spirals consistently featured elliptical rather than circular cross-sections (as would be anticipated for ideal superhelices) with mean ± s.d. major and minor axis lengths of 20.1 ± 5.6 nm and 8.6 ± 4.6 nm in the pointed-end directed force condition versus 14.0 ± 6.8 nm and 6.2 ± 3.1 nm in the barbed-end directed force condition (Fig. 2c,d). The major axes were randomly oriented relative to the plane of the ice film (Fig. S2d), suggesting this asymmetry results from F-actin’s helical architecture rather than flattening imposed by surface tension. We also occasionally observed sites featuring protruding densities suggestive of dislocated actin subunits (Fig. S2e), supporting an association between spiral formation and mechanical damage to F-actin that could produce filament ruptures. Collectively, these data show that myosin forces can directly generate F-actin domains featuring an asymmetric superhelical spiral morphology.

### Transient forces elicit superhelical F-actin

Myosin force generation is complex, featuring ATP hydrolysis-coupled F-actin binding and unbinding during the motor’s mechanochemical cycle^1,2^. To assess whether these features of myosin force generation could contribute to directionality-insensitive F-actin superhelix formation, we conducted coarse-grained molecular dynamics simulations (Fig. 2e; Methods; Fig. S2f,g; Video S6). Actin subunits were abstracted as spherical bodies arranged in a canonical F-actin lattice of four hundred protomers, linked by elastic bonds at inter-subunit interfaces. The filament (which lacks polarity in our simulations) was fixed at one end, then either tension (mimicking the pointed-end directed force condition) or compression (mimicking the barbed-end directed force condition) was applied.

The application of constant compression nearly instantaneously resulted in superhelix formation (Fig. 2f). The helical architecture of the filament transduces the compressive force into a torsional rearrangement that produces supercoiling. Conversely, the application of constant tension produced minor conformational fluctuations without striking architectural rearrangements (Fig. 2f). However, transient tension, mimicking motor unbinding after a powerstroke, elicited spiraling as the filament underwent elastic recoil (Fig. 2f). Transient compression conversely diminished spiraling upon recoil, although superhelical features nevertheless persisted after force was removed (Fig. 2f). We observe shorter wavelengths in transient compression vs. transient tension (Fig. 2g), consistent with our experimental observations of the barbed-end directed vs. pointed-end directed force conditions. Furthermore, transient tension and both compression conditions all produced superhelices with elliptical cross-sections (Fig. 2f), consistent with this being a feature which emerges from F-actin’s helical architecture. Collectively, these modelling studies suggest motor unbinding after a powerstroke is likely to be particularly important for the counterintuitive formation of superhelices in the pointed-end directed force condition. Myosin’s biophysical properties thus give rise to the directional insensitivity of superhelix formation, which is an intrinsic response of F-actin’s helical architecture to effective compression along the filament axis.

### Superhelical F-actin features distinct lattice architecture and subunit deformations

We next sought to visualize the structure of myosin force-evoked superhelical F-actin using single particle cryo-EM. We focused on the dual motor condition, as we found it to generate the greatest abundance of superhelices, collecting datasets in the absence and presence of ATP for detailed comparative analysis (Fig. S3, Table S1). We adapted a neural network-based approach we previously developed for analyzing F-actin bending^56^ to estimate the signed curvature of filament segments (Methods; Fig. S3a,b), facilitating specific detection of oscillatory domains for single particle analysis. The overall distribution of absolute curvatures in both datasets deviated markedly from that of F-actin undergoing thermally driven bending fluctuations^56,63^, with an increased proportion of low curvature segments, likely due to myosins tethering F-actin to the carbon film (Fig. S3c). We nevertheless observe a long tail of highly curved segments, which was significantly increased in the presence of ATP (Fig. S3c). We focused on filament regions featuring contiguous stretches of sharp positive and negative curvature (Methods). After extensive classification (Fig. S4a), a subset of 13,146 segments produced a consensus 9.5 Å resolution asymmetric reconstruction spanning 25 protomers (Fig. 3a; Fig. S4b; Fig. S5). Stitching multiple copies of this reconstruction (Fig. 3a) yielded a volume morphologically similar to superhelices observed in tomograms (Fig. 2a,b; Video S7). One notable discrepancy between this stitched reconstruction and the tomography and simulation results is the presence of a circular, rather than elliptical, cross-section. We infer this to be the consequence of averaging a heterogeneous set of oscillatory domain segments, each with potentially uniquely varying curvature. Consistently, cryoSPARC three-dimensional variability analysis (3DVA) supports extensive conformational variability (Fig. S4b), even within the relatively homogeneous particle subset isolated through classification. This heterogeneity, along with the low number of particles incorporated into the final reconstruction (an average of only 1 per 2.4 micrographs), limited the achievable resolution.

**Fig. 3:**
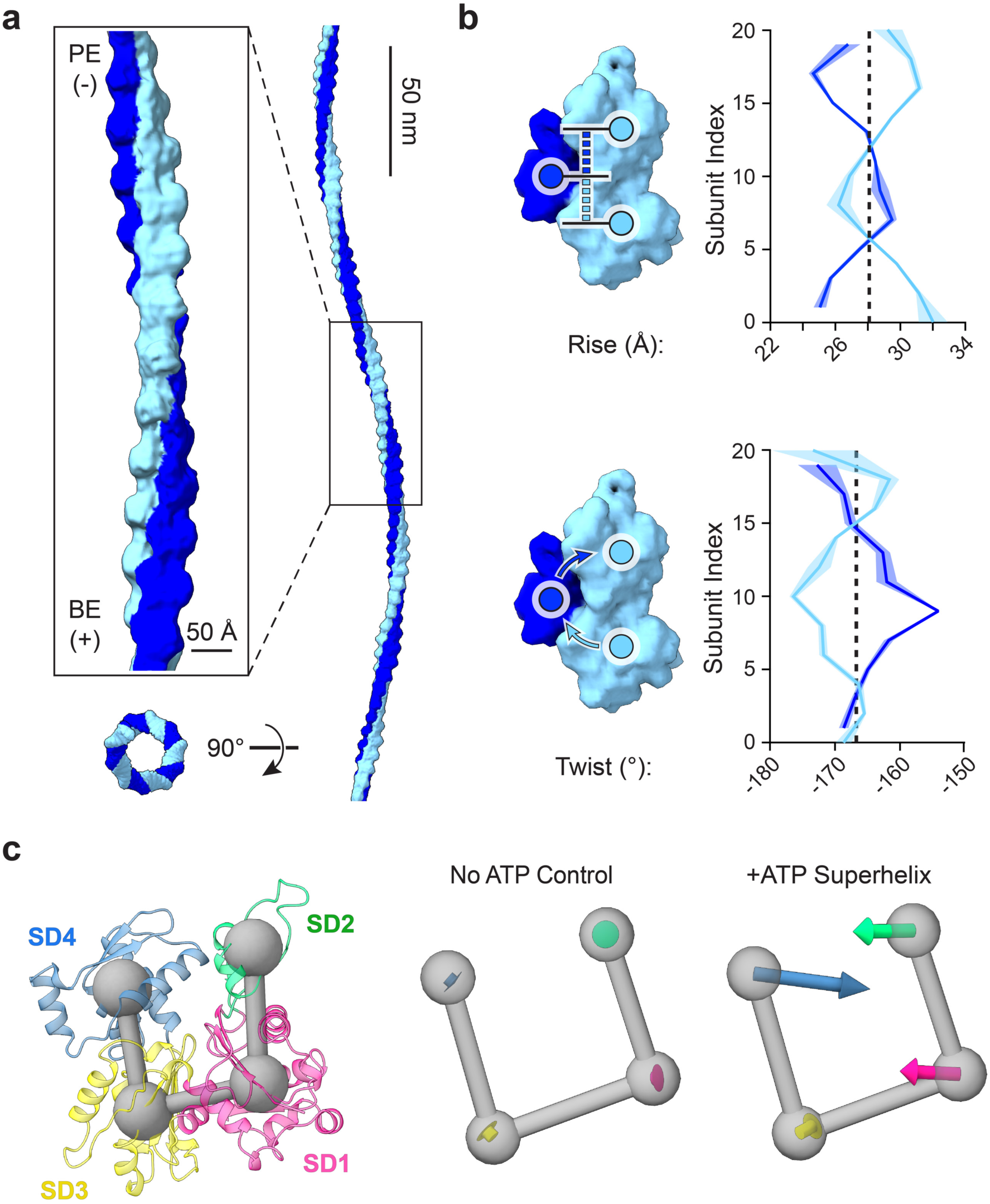
Superhelical F-actin features unique architectural remodeling and subunit deformations. **a**, Left: cryo-EM density map of superhelical F-actin reconstructed from the dual motor condition. Strands are colored in alternating shades of blue. BE: barbed end; PE: pointed end. Right: Stitch of five copies of the map, recapitulating the morphology of oscillatory domains. **b,** Diagrams (left) and plots (right) of instantaneous helical parameters. Strands are colored as in a. Shaded regions represent 95% CI from 3 independent analyses. Vertical dashed lines indicate parameters of canonical F-actin (ref. 56). **c**, Left: ribbon diagram of an actin subunit, with subdomains colored in varying shades. Subdomain centroids are indicated by connected gray spheres. Right: averaged subdomain displacements (scaled 15X for visualization) after MDFF analysis relative to a canonical F-actin subunit (PDB 8D13) for the indicated conditions.

To probe changes in F-actin’s helical lattice, we rigid-body docked actin subunits into the map and analyzed their positioning along a deformed filament axis determined by numerical fitting (Methods; Fig. 3b; Fig. S5). We also performed this analysis on two control reconstructions determined from the -ATP dataset (Fig. S5a,e), which featured low curvature and helical parameters correspondingly highly similar to those of canonical F-actin (Fig. S5e). Conversely, in the superhelical F-actin reconstruction, we observe extensive architectural remodeling (Fig. 3b). Helical twist is altered in a pattern similar to that we previously reported for bending deformations^56^ (Fig. S5f), where one strand is over-twisted and the other under-twisted in a wavelike pattern, while maintaining the canonical F-actin twist as their instantaneous average. This supports twist-bend coupling as a general response of the F-actin lattice to the introduction of curvature^64^. However, superhelical F-actin also features distinct and extensive modulation of helical rise (Fig. 3b), which is only modestly impacted by bending^56^ (Fig. S5f). The strands feature alternating over-extension and under-extension, once again occurring in a wavelike pattern around an instantaneous average matching that of canonical F-actin (Fig. 3b).

We next examined structural deformations at the level of individual actin protomers (Fig. 3c; Fig. S6). Due to the modest resolution of the map, we performed molecular dynamics flexible fitting in ISOLDE with extensive restraints that restricted changes to the relative repositioning of actin’s four subdomains (Methods). While fitting in the control reconstructions uncovered negligible transitions (Fig. 3c; Fig. S6), superhelical F-actin featured substantial subunit deformations characterized by an inward compression of subdomains 1 and 4 into the nucleotide cleft. This contrasts with bent F-actin, which we previously found features subunit shearing outwards from the filament axis^56^. In addition to superhelical F-actin featuring specific lattice architectural remodeling and subunit deformations, the absolute curvature distribution of segments which met our oscillating curvature selection criteria (Fig. S3c) deviates markedly from that previously observed for bending fluctuations^56^. These data collectively suggest that myosin force-evoked superhelical F-actin is structurally distinct from F-actin undergoing bending deformations.

### α-catenin detects and modulates the myosin-force evoked F-actin structural landscape

As myosin forces have been shown to regulate F-actin engagement by force-sensitive ABPs^23,34,38,39^, we postulated that these proteins may discriminate features of superhelical F-actin through binding contacts. To test this hypothesis, we focused on the isolated actin-binding domain (ABD) of the cell-cell adhesion protein αE-catenin, which displays enhanced F-actin binding in the presence of myosin forces^34^. We prepared dual motor cryo-EM specimens with 1 µM α-catenin and 0.6 µM F-actin (Methods), a sub-saturating regime of α-catenin where force-sensitive binding is observed. After multiple rounds of classification (Fig. S7), we obtained a 10.4 Å resolution consensus three-dimensional reconstruction (Fig. S8; Table S1).This map features asymmetric F-actin decoration, with α-catenin preferentially binding along one strand, then switching to the other at the crossover where the strands exchange sides along the filament. This binding mode is distinct from previously reported structures obtained under saturation binding conditions in the absence of force, where symmetric binding along both strands was observed^34,65^.

To probe the links between F-actin conformational remodeling and force-dependent α-catenin binding, we performed three-dimensional variability analysis (3DVA; Methods; Fig. S7; Video S8). Inspection of the first variability component revealed variations in α-catenin binding that correlated with F-actin rearrangements (Fig. 4a). To quantify α-catenin binding, we rigid-body docked the high-resolution structure of one complete α-catenin ABD-F-actin interface (PDB 6UPV^34^), composed of 2 actin protomers and one ABD, at each binding position in each frame of the 3DVA trajectory. We then measured the integrated intensity of the map within a mask calculated from the ABD coordinates, a proxy for α-catenin occupancy. When examining filament curvature (Methods) versus the average α-catenin intensity of each frame, we observed two regions in the trajectory featuring high curvature (Fig. 4b), only one of which also displays high average α-catenin intensity. This suggests α-catenin engages a specific mechanically- excited conformational regime rather than globally recognizing deformed F-actin. We next analyzed the helical parameters of the 3DVA frames in these two trajectory regions (Fig. 4c; Fig. S9a). While the low α- catenin intensity frames are similar to unbound superhelical F-actin, the high α-catenin intensity frames display a distinct pattern, featuring even larger rise deviations that persist for a greater distance along each strand without oscillations, coupled to suppression of twist deviations.

**Fig. 4:**
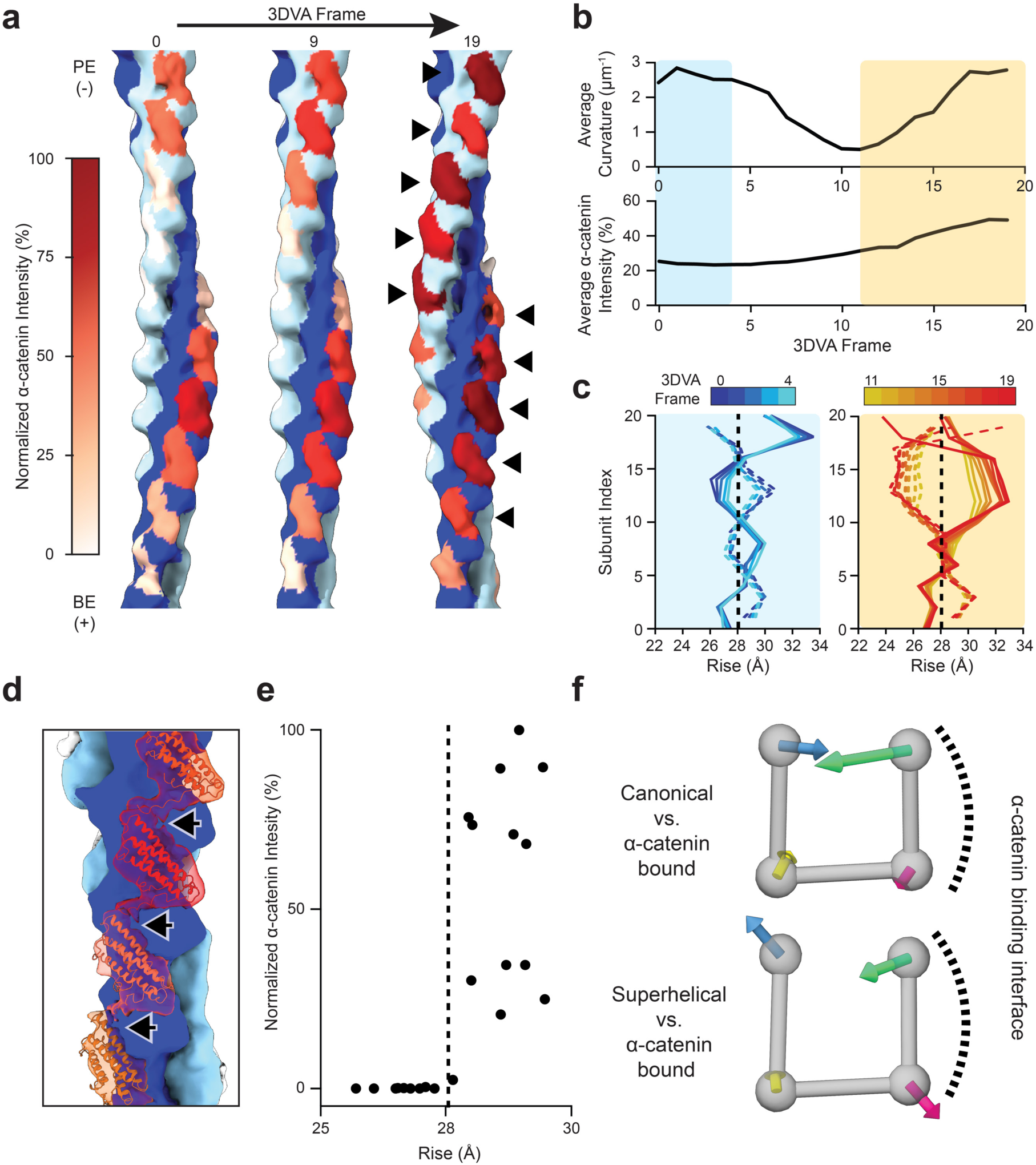
α-catenin detects and reciprocally modulates force-evoked structural changes in F-actin. **a**, Frames from 3DVA variability analysis of the α-catenin ABD–F-actin complex reconstructed in the dual motor force condition, highlighting alternating binding along F-actin strands (arrowheads) and varying intensity of α-catenin density across the trajectory**. b,** Quantification of average filament curvature (top) and average α-catenin intensity (bottom) across the 3DVA trajectory. Boxes indicate regions with high filament curvature and either low (blue) or high (orange) α-catenin intensity. **c**, Plots of instantaneous helical rise from trajectory regions indicated in b. Vertical dashed lines indicate canonical F-actin rise. **d**, Docking analysis (PDB: 6UPV) of consensus reconstruction highlights inter-ABD contacts mediated by α- catenin’s C-terminal extension (arrows). **e**, Quantification of α-catenin intensity versus instantaneous rise of the consensus reconstruction (Fig. S9b). Vertical dashed line indicates canonical F-actin rise. **f**, Averaged subdomain displacements (scaled 15X for visualization) after MDFF analysis of the consensus map versus canonical F-actin (top, PDB: 8D13) and superhelical F-actin (bottom, MDFF model from Fig. 3f).

To examine how these F-actin architectural changes are linked to α-catenin binding, we selected particles which contributed to the high α-catenin intensity frames. Refinement produced a 12.3 Å resolution reconstruction (Fig. S9b; Fig. S8; Table S1) featuring a similar pattern of α-catenin binding (Fig. S9c) and helical parameter modulation (Fig. S9d). Despite the moderate resolution of the map, density connecting longitudinally-adjacent ABDs was clearly visible (Fig. 4d). Docking analysis assigns this density to the ordered portion of α-catenin’s C-terminal extension (residues 865-871), a region necessary for force-activated actin binding that has previously been implicated in mediating inter-ABD contacts^34^. We then examined the relationship between α-catenin intensity and the instantaneous rise at each protomer index, which revealed preferential α-catenin engagement at positions featuring rise greater than that of canonical F-actin (Fig. 4e).

To assess whether these high α-catenin intensity binding positions are associated with specific actin subunit remodeling, we performed MDFF and examined actin subdomain rearrangements (Fig. 4f; Fig. S9e). Regardless of α-catenin intensity, all of the actin subunits captured in our reconstruction feature repositioning of subdomain 2, a flexible segment of the protein that mediates inter-subunit contacts (Fig. 4f; Fig. S9e). This observation suggests substoichiometric α-catenin binding to mechanically-excited F-actin also modulates the conformation of actin subunits at unbound positions, consistent with subdomain 2’s previously established role in facilitating F-actin structural plasticity^46,52–56^. However, high-occupancy positions feature a characteristic actin subunit rearrangement specifically adjacent to direct binding contacts with the α-catenin ABD, where subdomain 2 is pulled away from the filament core (Fig. S9f). This produces an actin subunit conformation distinct from both canonical F-actin and unbound superhelical F-actin (Fig. 4f; Fig. S9e). Notably, this subdomain 2 rearrangement is coupled to a reduction in the compression of subdomains 1 and 4 into the nucleotide cleft, which is likely to relieve steric strain on the subunit. Taken together, these data show α-catenin asymmetrically binds F-actin in the presence of myosin forces, preferentially engaging positions with extended rise while suppressing twist deviations. These lattice transitions are likely stabilized by cooperative inter-α-catenin ABD contacts, as well as actin subdomain 2 rearrangements that relieve force-induced steric strain on the subunits composing the remodeled filament lattice.

## Discussion

We find that myosin activity can generate superhelical domains in actin filaments *in vitro*, which are also present at load-bearing cytoskeleton-adhesion interfaces in cells. While micron-scale F-actin superhelices have previously been observed in modified gliding assays^66^, to our knowledge nanoscale rearrangements such as those we report here have not previously been described. We are aware of one previous report of F-actin featuring oscillatory nanoscale curvature, which was observed with cryo-EM under supraphysiological conditions of saturation myosin binding^67^. In this kinetic trapping study, photo-uncaging of ATP transiently induced filament curvature prior to myosin dissociation, likely through direct allosteric effects at actin-myosin interfaces as motors stochastically bound nucleotide. Our work shows that myosin force generation can elicit spiraling in filament regions distal from motor binding sites, a mechanism for generating a specific F-actin conformational mark that is accessible to force-sensitive ABPs (Fig. 5). Although the precise biophysical mechanism for superhelical domain formation remains to be determined, our simulations suggest that cycles of myosin applying force and unbinding elicit a characteristic rearrangement due to F-actin’s helical architecture. Future work will be required to analyze the precise force regimes and timescales associated with generating superhelical F-actin, as well as whether its formation is reversible or invariably leads to filament rupture.

**Fig. 5:**
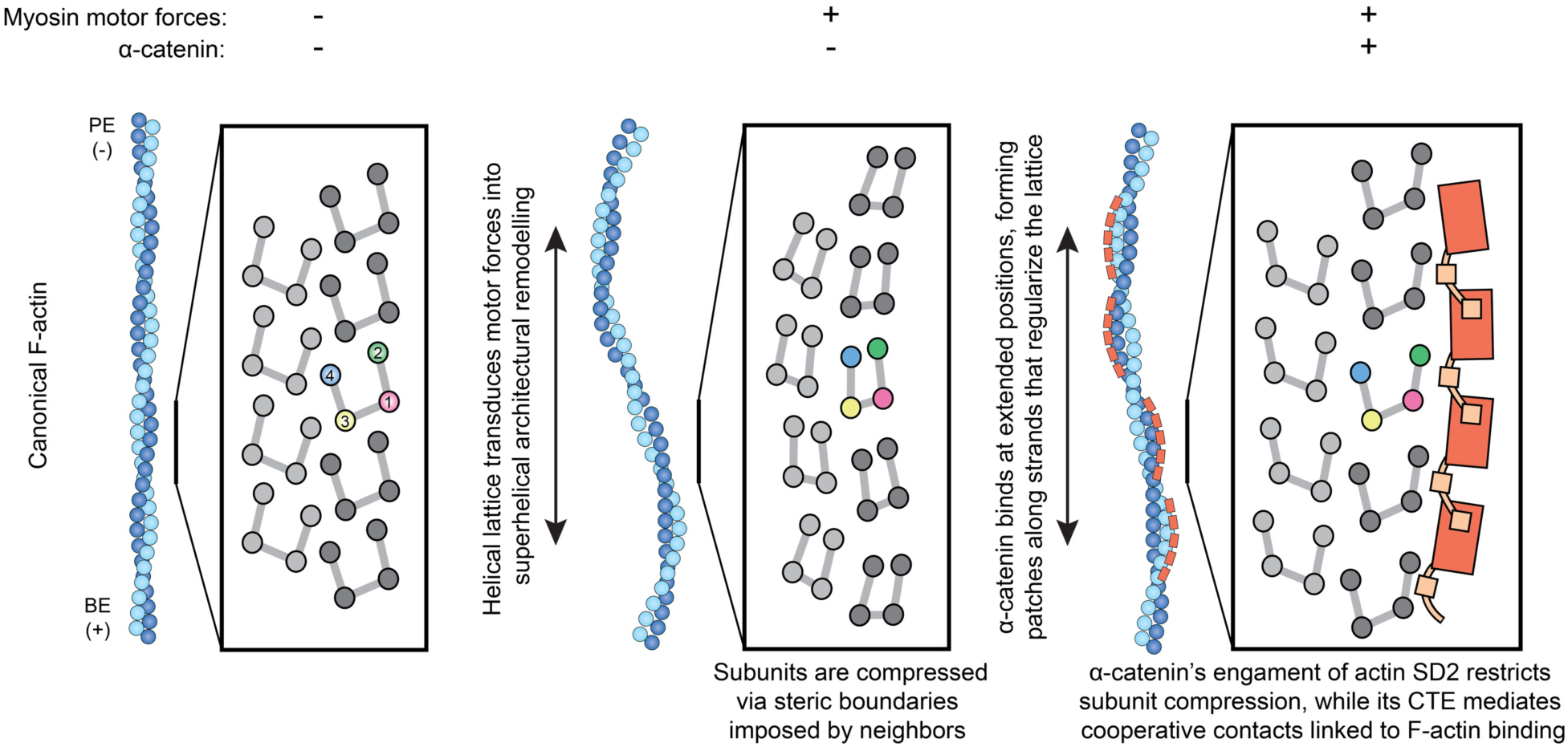
Conceptual model of myosin force transduction through mechanosensitive F-actin binding. Cartoon schematizing generation of destabilized superhelical F-actin by myosin forces, which is detected and stabilized through cooperative binding contacts by α-catenin.

While eukaryotic actins are highly conserved in sequence and structure, divergent prokaryotic actin homologs with diverse filament architectures have been reported^68,69^, including the formation of a stable superhelix by *Bacillus thuringiensis* parM^70^. In eurkaryotes, the actin fold’s latent potential for forming superhelical assemblies may have been specifically harnessed as a mechanism for transducing myosin forces into downstream biochemical processes through differential ABP engagement, concomitant with the unique appearance of cytoskeletal motors in this lineage^71^. Selection for the capacity to undergo such mechanically-evoked transitions could provide one explanation for the extreme sequence conservation of eukaryotic actins despite the divergence of ABPs across species, consistent with prior speculation^15^.

Myosin force-evoked superhelical F-actin features specific lattice architectural remodeling and actin protomer deformations that are distinct from those elicited by bending forces^56^, which could facilitate the discrimination of these two force regimes by force-sensitive actin-binding proteins (Fig. 5). We find that the α-catenin ABD preferentially engages positions featuring extended rise, a hallmark of superhelical F-actin. This asymmetric binding further modulates F-actin’s mechanically-excited conformational landscape, accentuating rise changes while regularizing twist. As superhelices are likely to be associated with mechanical damage (Fig. S2e), we speculate α-catenin binding stabilizes F-actin in the presence of myosin forces, which could enhance adhesion-cytoskeleton coupling in the presence of contractility.

Mechanistically, α-catenin’s F-actin binding contacts mediate repositioning of actin subdomain 2. In addition to α-catenin providing stabilizing binding energy, this repositioning can be explained by bound α-catenin modifying the space available for actin subunits to occupy as they are deformed by lattice rearrangements, consistent with a steric boundaries framework for mechanical regulation of F-actin^56^. Furthermore, our structural data suggest inter-ABD cooperative binding interactions mediated by α- catenin’s C-terminal extension support its accumulation along individual strands, rationalizing the requirement of this region for force-activated actin binding^34^. Other force-sensitive actin binding proteins feature tandem arrays of LIM domains with precise interdomain linker lengths^38,39^, suggesting they may also recognize an F-actin structural signature spanning multiple subunits. We speculate multi-actin subunit engagement mechanisms confer sensitivity to the Ångström-scale rearrangements that occur within individual actin promoters and at the interfaces between them. These subtle structural transitions accumulate into substantial lattice architectural changes that can be more readily discriminated through binding contacts at the multi-subunit scale. Further structural characterization of force-activated F-actin complexes will be necessary to probe the generality of this concept, which should be accessible through the approaches we introduce here. More broadly, we anticipate the capacity to directly visualize these active mechanical signaling assemblies will advance our mechanistic understanding of force transduction pathways and their malfunction in disease.

## Acknowledgements

We gratefully acknowledge Jenny Hinshaw (NIDDK) for use of her F20 electron microscope, and Clare Waterman (NHLBI) for use of her epifluorescence microscope. We also thank Johanna Sotiris, Honkit Ng, and Mark Ebrahim at the Rockefeller University Evelyn Gruss Lipper Cryo-EM Resource Center (CEMRC) for their assistance with cryo-fluorescence, cryo-ET, and cryo-EM data collection. A.G.C. was supported by NIH T32 GM115327, and D.Y.Z.P. was supported by NIH / NHLBI fellowship F31HL165906. P.G. was supported by a Rockefeller University Women in Science Postdoctoral Fellowship., and K.H. was supported by a Rockefeller University Anderson Center for Cancer Research Postdoctoral Fellowship. A.J.N was supported by NIH / NIGMS fellowship F32GM128303. This work was funded by grants from the NIH (R01GM141044 and 5DP5OD017885), the Alfred P. Sloan Foundation (G-2020- 14047), and the Pew Charitable Trusts to G.M.A., and NIH grant HL004232 to J.R.S. This research was also supported by the Stavros Niarchos Foundation (SNF) as part of its grant to the SNF Institute for Global Infectious Disease Research at the Rockefeller University.

## Author contributions

D.Y.Z.P. prepared and imaged cellular cryo-ET specimens, which were computationally processed by K.H.; P.S.G. initially developed the myosin reconstitution assay for cryo-EM studies with assistance from Y.T. and J.R.S.; A.G.C. and P.S.G. prepared and imaged myosin reconstitution specimens, and L.M. prepared α-catenin specimens, which were imaged by A.G.C.; A.G.C. and M.J.R. performed cryo-EM analysis and data interpretation, using computational tools developed by M.J.R.; X.S. performed coarse-grained molecular dynamics simulations; A.G.C. performed *in vitro* cryo-ET studies with assistance from A.J.N., which were analyzed by M.J.R.; A.G.C., M.J.R., and G.M.A. analyzed data and wrote the paper with input from all authors; G.M.A. supervised the study and conceived of the project.

## Data Availability

Cryo-EM density maps have been deposited in the EMDB with the following accession codes: Myosin force-evoked superhelical F-actin (EMD-46426); Control myosin tethered F-actin -ATP 1 (EMD-46427); Control myosin tethered F-actin -ATP 2 (EMD-46428); Consensus force-activated α-catenin–F-actin complex (EMD-46429); 3DVA sorted force-activated α-catenin–F-actin complex (EMD-46431). Additional raw and processed data files are available at Zenodo. Trained neural networks and denoised *in vitro* tomograms with filament traces are available at https://doi.org/10.5281/zenodo.12702199. Trained neural networks used for single particle analysis particle picking, flexibly fit PDB models, and variability analysis maps and models used for data analysis are available at https://doi.org/10.5281/zenodo.12702799.

## Resource Availability

All reagents and resources reported in this study are freely available from the corresponding author. Requests should be directed to Gregory M. Alushin.

## Code Availability

Custom code reported in this study is available at https://github.com/alushinlab/squiggle as open source.

---

**Fig. S1:**
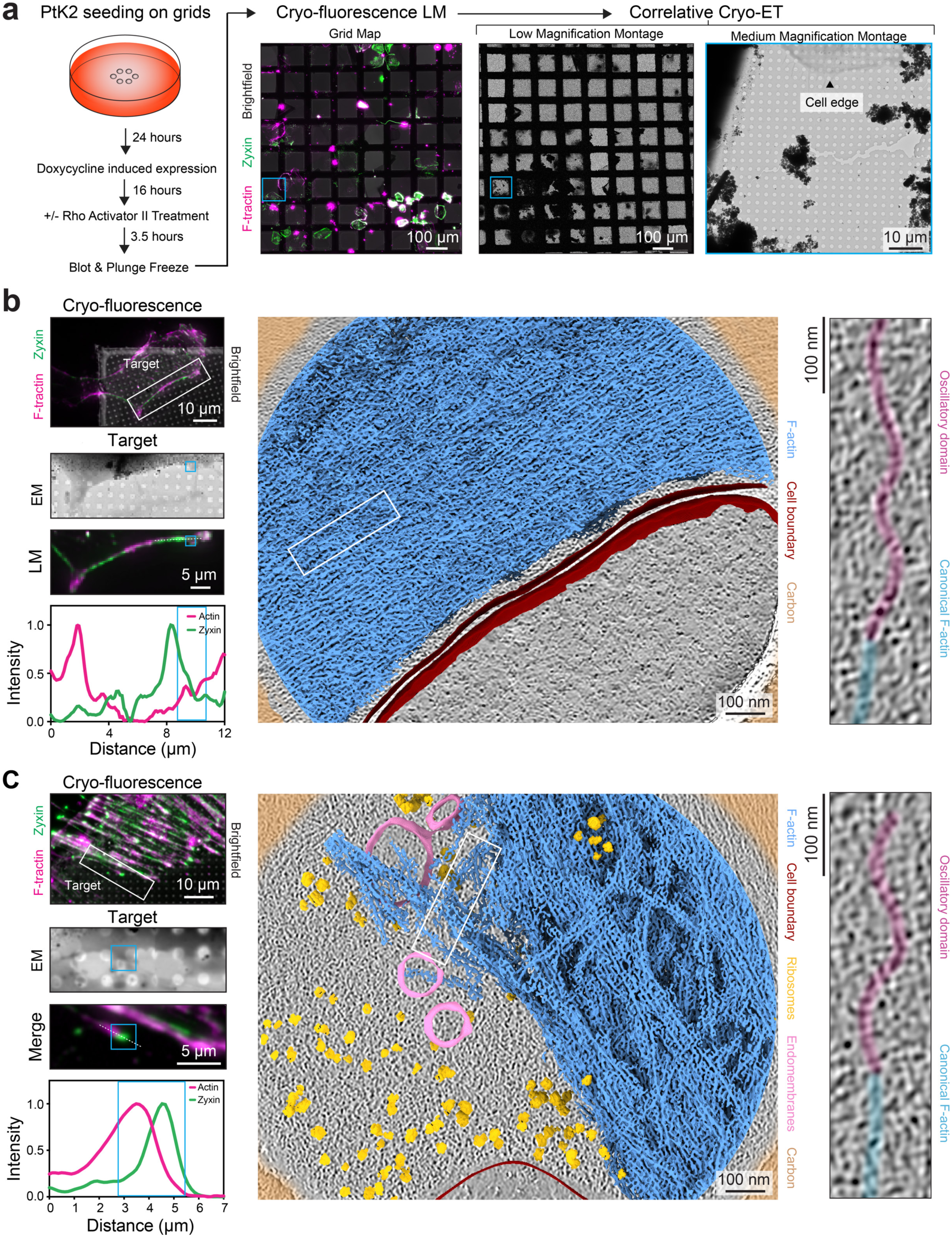
Correlative cryo-fluorescence and cryo-electron tomography. **a**, Schematic of sample preparation and data collection workflow. **b and c**, Additional examples of oscillatory domains imaged in zyxin-enriched adhesion sites from N = 2 independent experiments. The site in **b** features parallel bundled F-actin and was visualized in a cell not treated with Rho Activator II, while the site in **c** features more disorganized F-actin and was visualized in a cell treated with Rho Activator II. Correlative LM / EM and tomograms / segmentations are presented as in Fig. 1a,b.

**Fig. S2:**
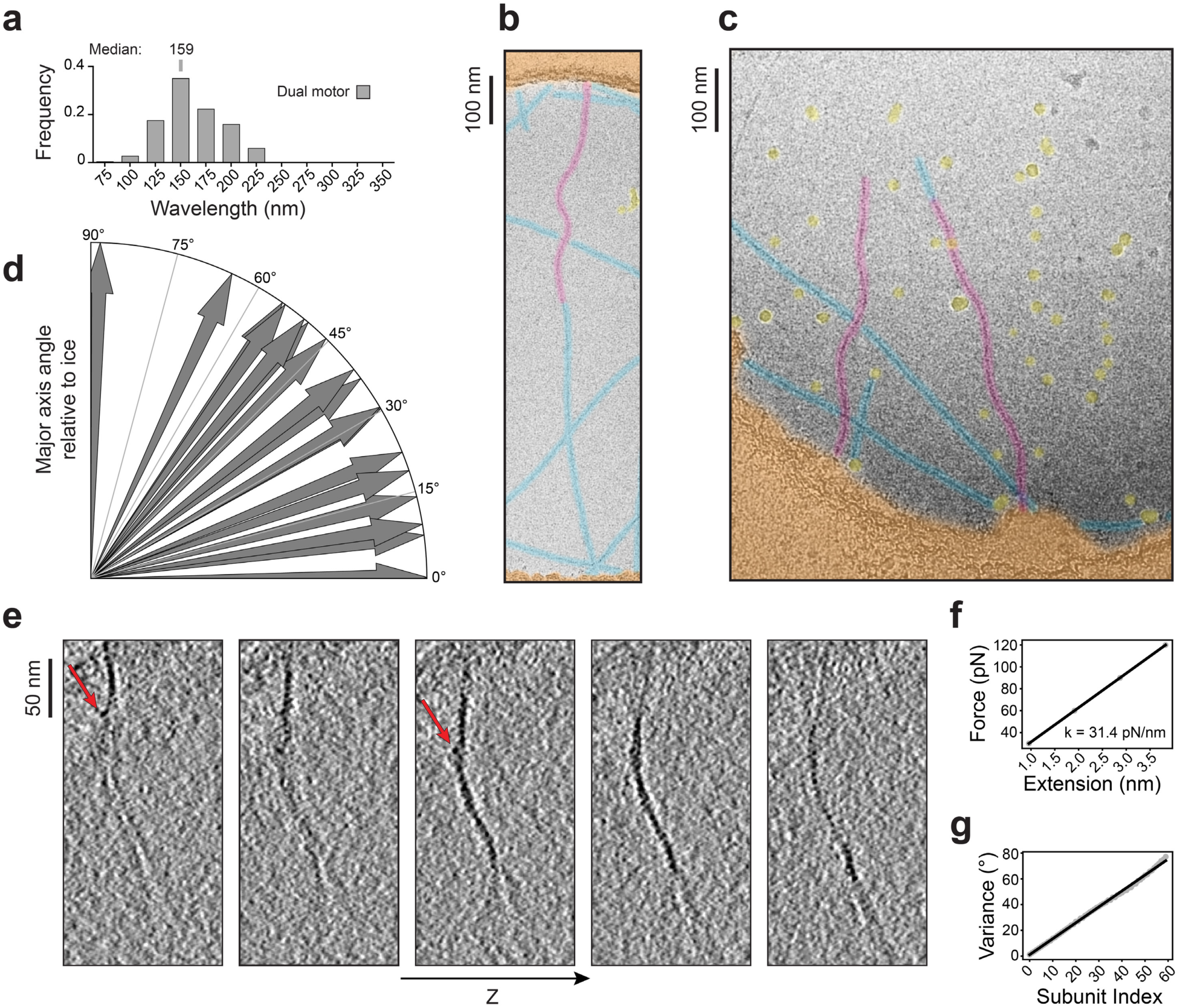
Analysis of F-actin oscillatory domain morphology. **a**, Quantification of oscillatory domain wavelengths in the dual motor condition. n = 251 from N = 3 independent experiments. **b and c**, False-colored cryo-EM images of the dual motor condition in the presence of ATP, displaying oscillatory domains in a hole-spanning filament (**b**) and in a pair of broken filaments (**c**). Oscillatory domains, magenta; canonical F-actin, blue; carbon film, orange; ice contamination, yellow. **d**, Polar arrow plot of oscillatory domain major axis orientation relative to the ice plane from n = 16 observations, where 0° corresponds to the major axis being parallel to the ice plane. **e**, Serial Z slices through an oscillatory domain tomogram (barbed-end directed force condition) featuring protruding densities consistent with subunit dislocations (red arrows). **f**, Calibration of the F-actin force-extension curve to determine harmonic bond stiffness in coarse-grained molecular dynamics simulations. **g**, Calibration of cumulative twist variance to determine the bending constants of the harmonic angle potential and the dihedral potential used in coarse-grained molecular dynamics simulations.

**Fig. S3:**
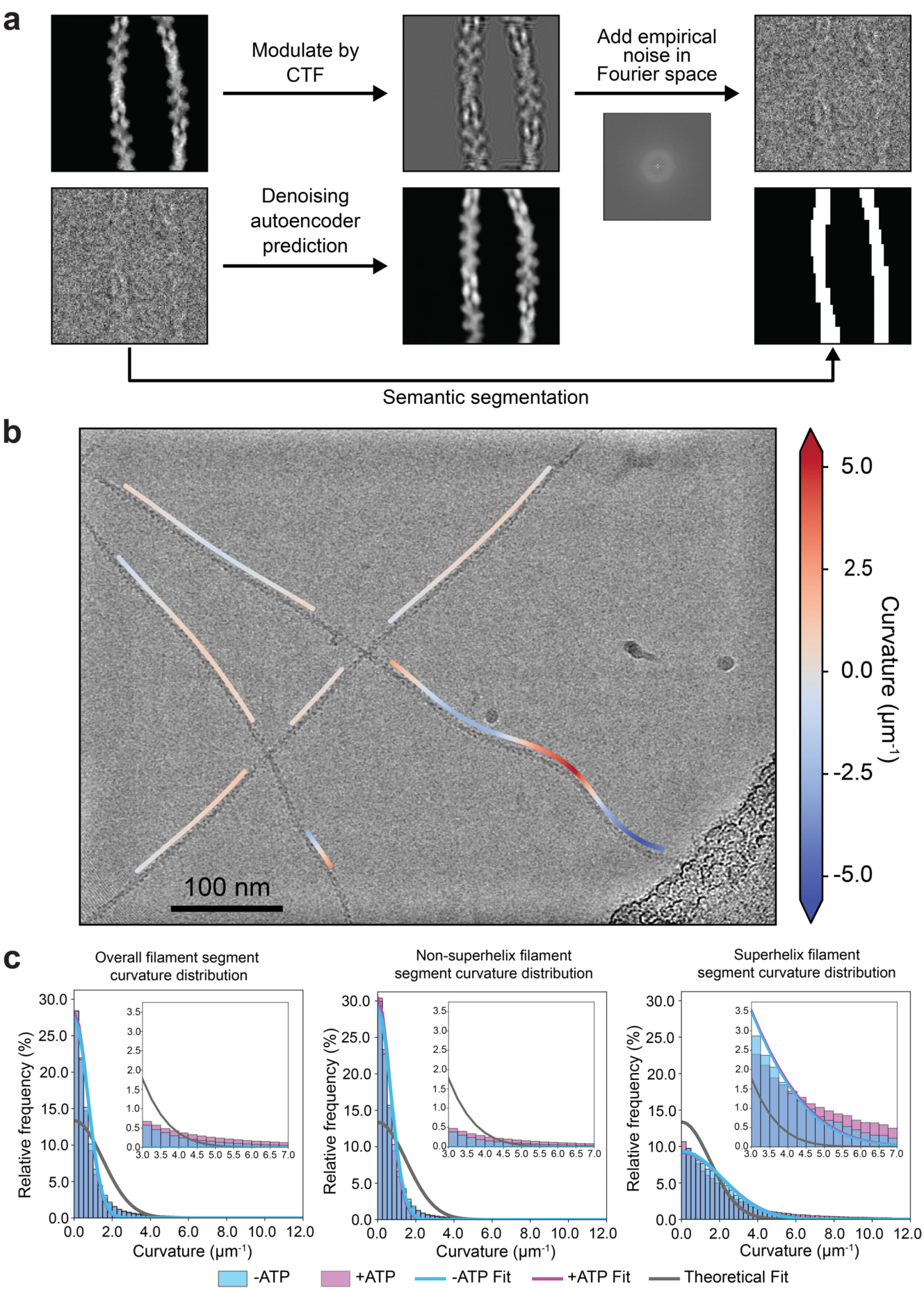
Adaptation of neural network picker and filament curvature analysis. **a**, Workflow for network training and subsequent picking. A synthetic particle dataset is generated by projecting PDB models of filaments featuring different computationally-generated curvatures, which are modulated by the CTF followed by the addition of a pink noise box (top). The network is then trained to denoise these noisy particles, which is then used as an input for semantic segmentation. After training with synthetic data, the network can be used to denoise and segment real data. **b**, Sample micrograph containing both straight filaments and filaments featuring oscillating curvature. Filaments are assigned estimated signed curvature values and categorized for subsequent selection. Traces are offset by one F-actin width for visualization. **c**, Curvature distributions of filaments identified in the dual myosin-motor evoked force condition (pink) or the –ATP control (blue). Measured curvature distributions of all picked filament segments (left), non-superhelical segments (middle), and superhelical segments (right) are shown as histograms, and modeled thermal bending fluctuation distributions are overlaid as blue, pink, and grey curves. The gray curve corresponds to a simple bending model described by equations 5-7 (Methods) using actin’s experimentally determined persistence length of 9 μm (ref. 63), while the pink and blue curves correspond to a model fitted with a multiplicative adjustment factor as described in detail in the Methods.

**Fig. S4:**
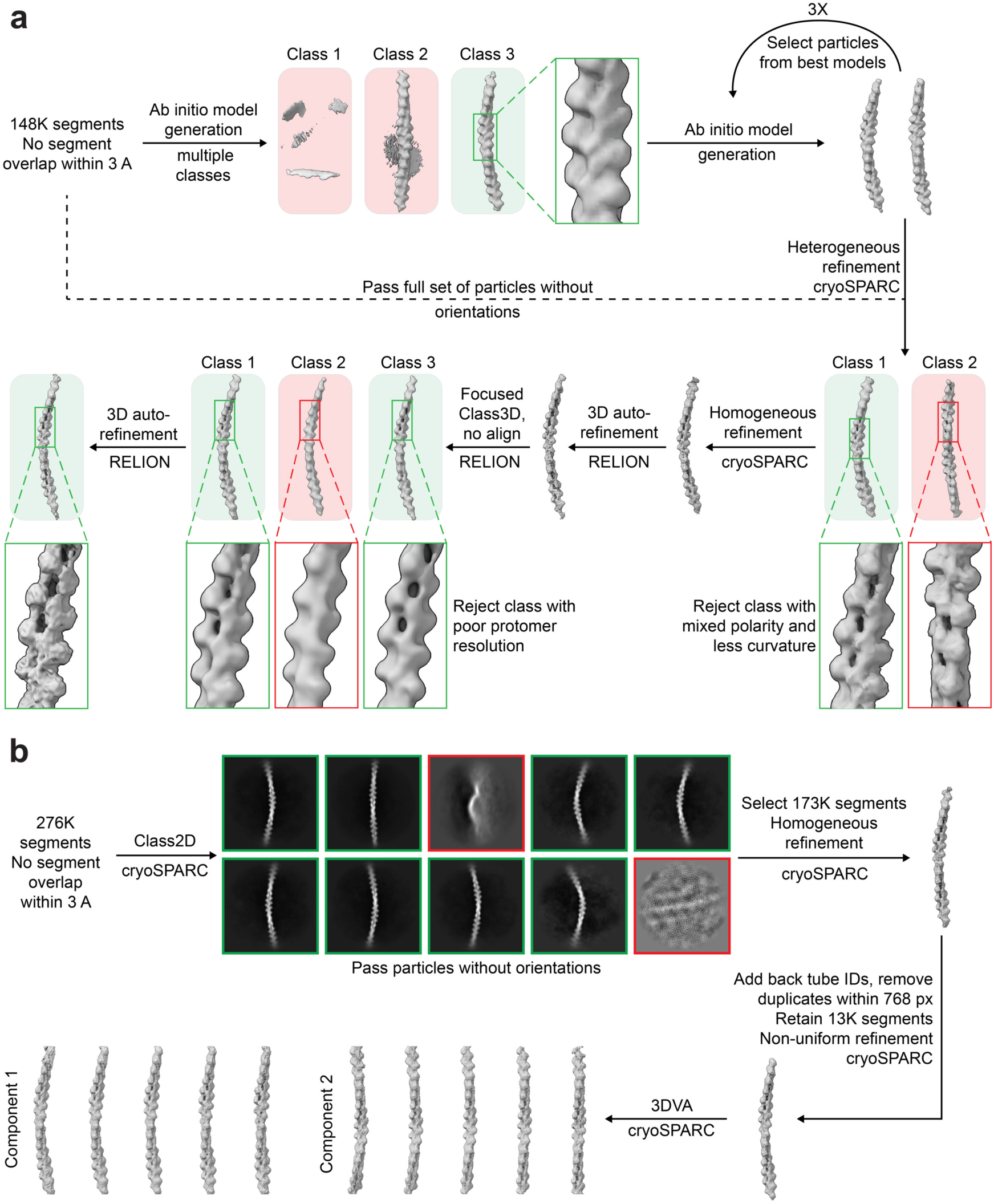
Superhelix structure determination workflow and variability analysis. **a**, Initial single particle cryo-EM data processing workflow for superhelical F-actin, using a single dataset collected for the dual motor condition. Transparent red and green boxes indicate rejected and accepted classes, respectively. **b**, Final processing workflow, incorporating data from two additional datasets to boost particle number and enhance the quality of the final map. Additionally, 3DVA variability analysis is displayed, highlighting the presence of continuous structural variability despite extensive classification.

**Fig. S5:**
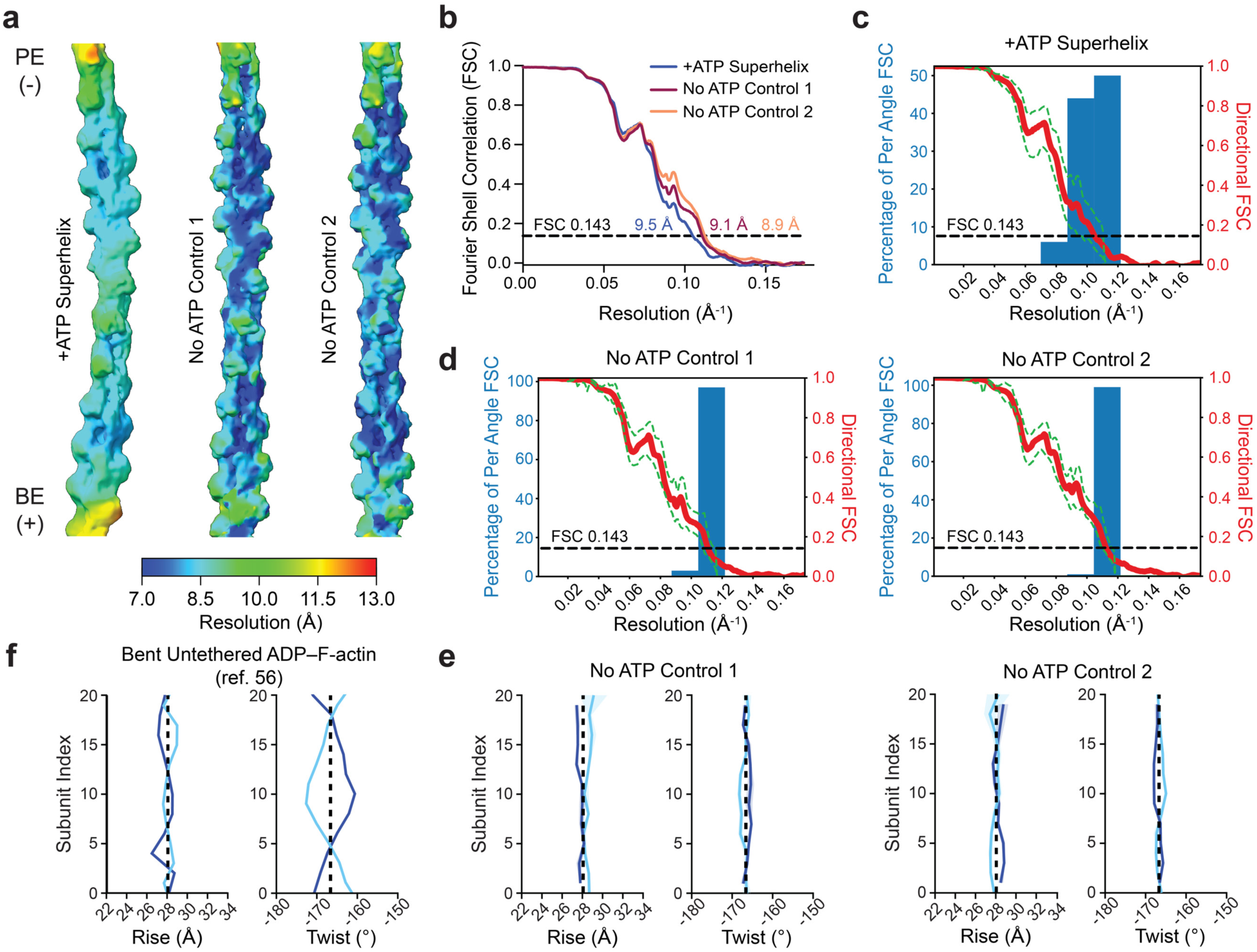
Resolution assessment and helical parameter analysis of superhelical and control F-actin. **a**, Local resolution of +ATP myosin-force evoked superhelical F-actin (left) and two independently reconstructed –ATP control maps (right). BE: barbed end; PE: pointed end. **b**, Global Fourier Shell Correlation (FSC) curves. **c and d**, 3DFSC curves for +ATP superhelical F-actin (**c**) and –ATP control reconstructions (**d**). **e**, Instantaneous helical parameters of –ATP control reconstructions, colored as in Fig. 3b. Shaded regions represent 95% CI from 3 independent analyses. **f**, Instantaneous helical parameters of bent, untethered ADP–F-actin with 5.4 μm^-1^ curvature, from ref. 56. Vertical dashed lines indicate parameters of canonical F-actin (ref. 56).

**Fig. S6:**
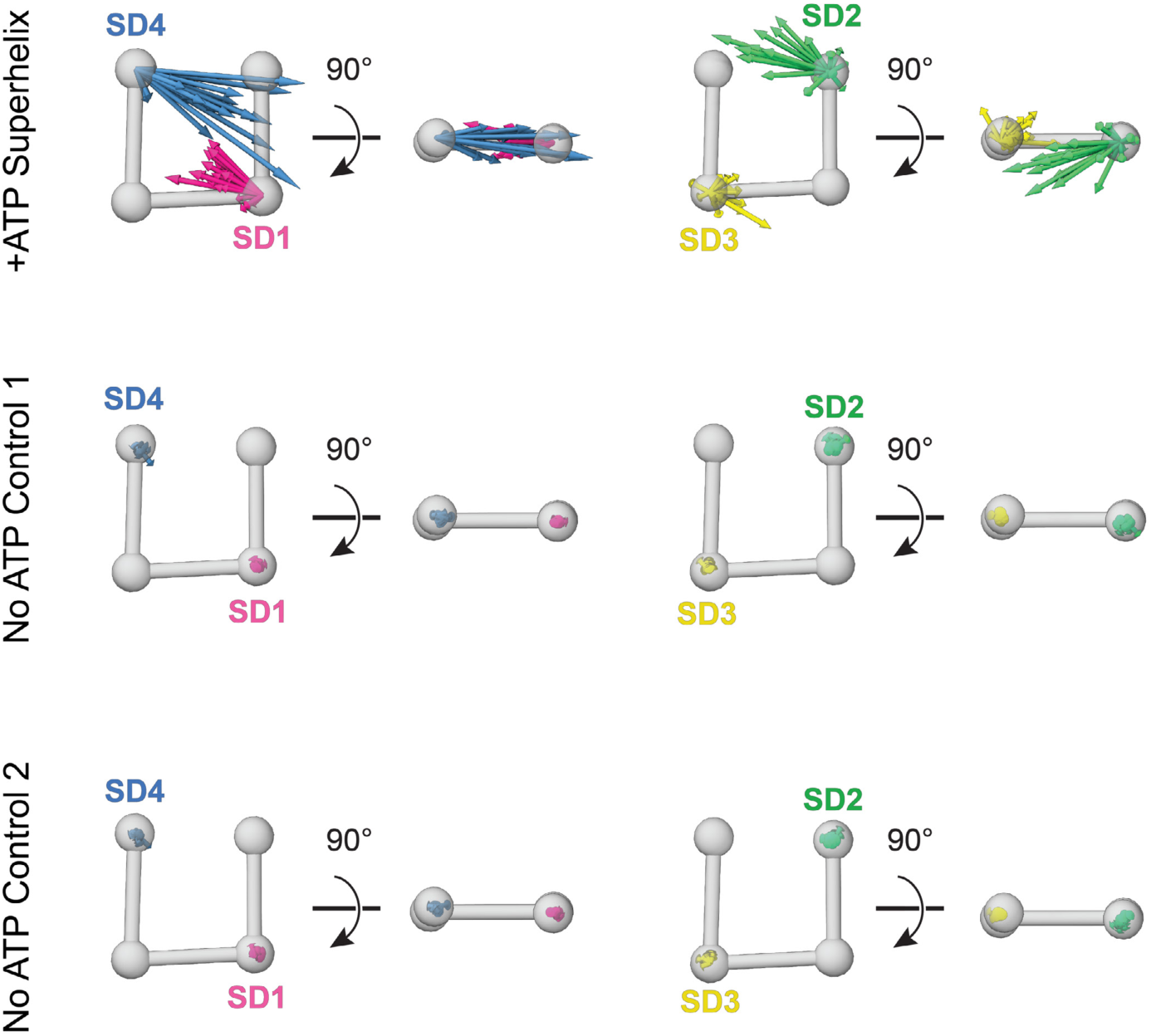
Subdomain displacements in superhelical F-actin. Superimposed subdomain displacement vectors from all protomers after MDFF analysis of +ATP myosin force-evoked superhelical F-actin (top) and –ATP control reconstructions (bottom). Subdomains 1 and 4 versus 2 and 3 are displayed separately for clarity, and vectors are scaled 15X for visualization. The averages of these vectors are displayed in Fig. 3c.

**Fig. S7:**
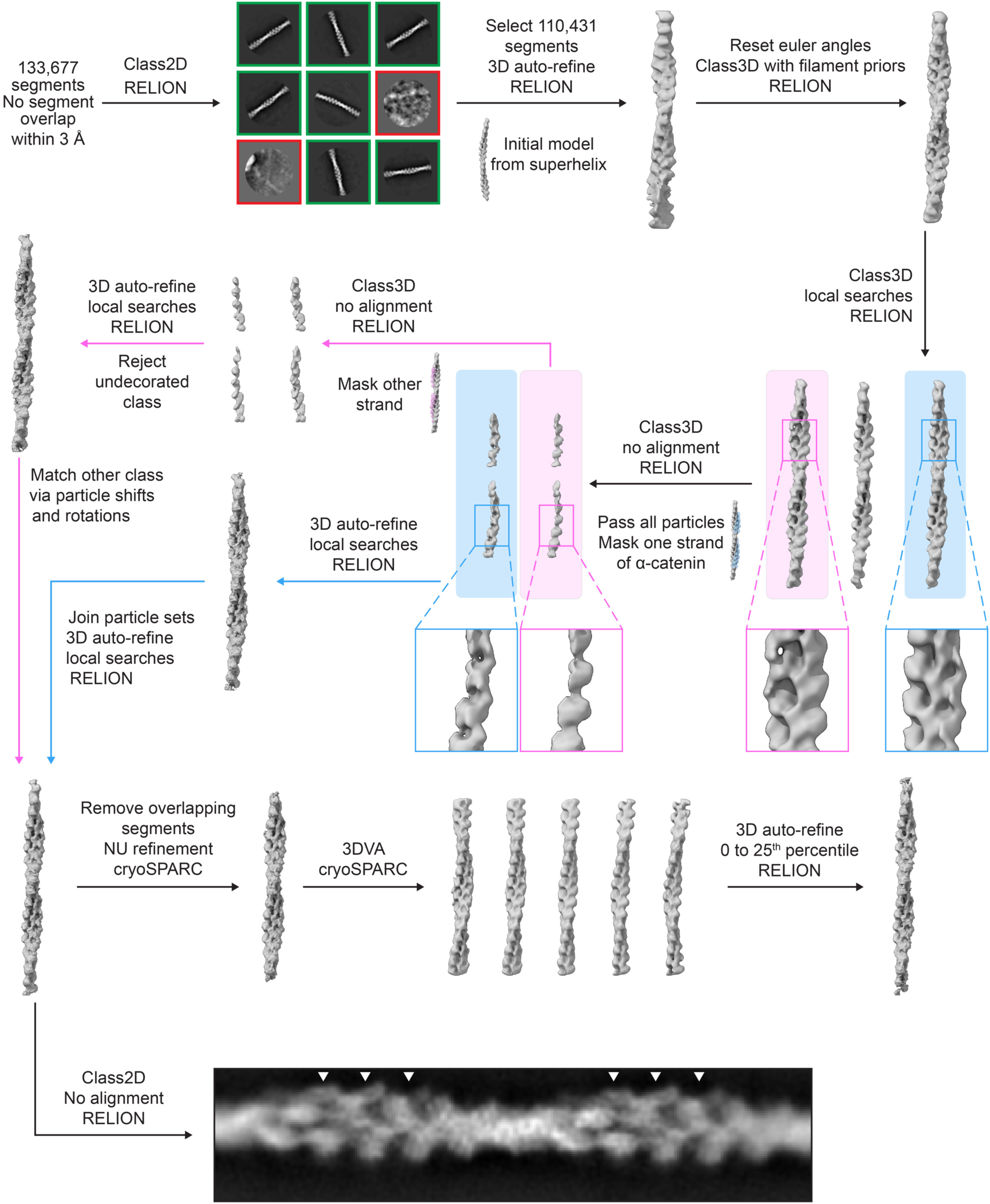
Cryo-EM processing workflow for the myosin force-activated *α*-catenin–F-actin complex. Top: Cryo-EM processing workflow for visualizing the force-activated *α*-catenin-F-actin complex, from a specimen prepared in the dual motor condition. Green and red boxes represent 2D class averages which were selected and rejected for additional processing, respectively. Bottom: a magnified 2D class average is displayed, highlighting preferential binding of *α*-catenin along one side of the filament (arrowheads) on alternating F-actin strands.

**Fig. S8:**
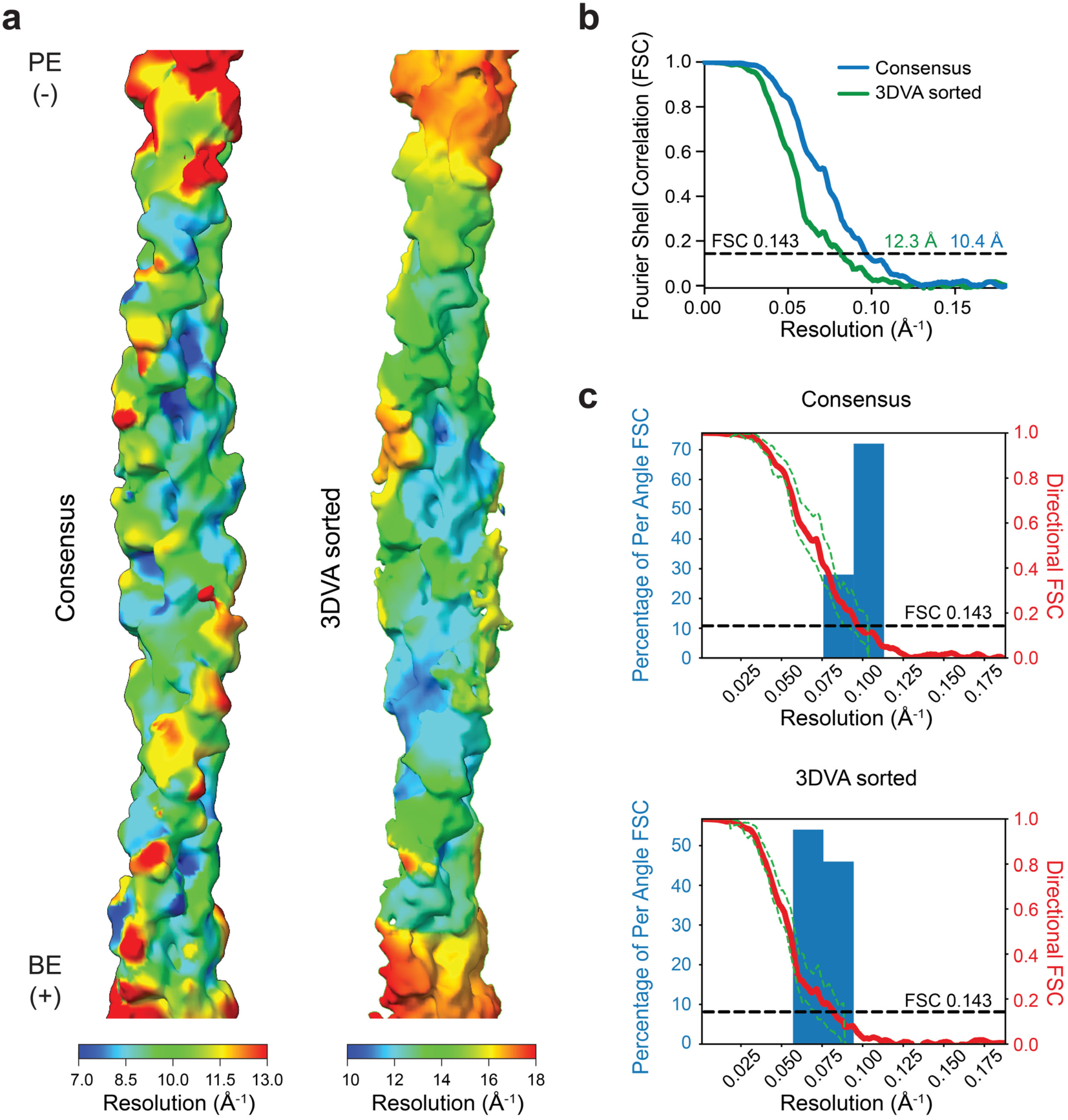
Resolution assessment of force-activated *α*-catenin–F-actin complex reconstruction. **a**, Local resolution of final consensus (left) and 3DVA sorted (right) *α*-catenin–F-actin complex reconstructions. BE: barbed end; PE: pointed end. **b**, Global FSC curves. **c**, 3DFSC curves.

**Fig. S9:**
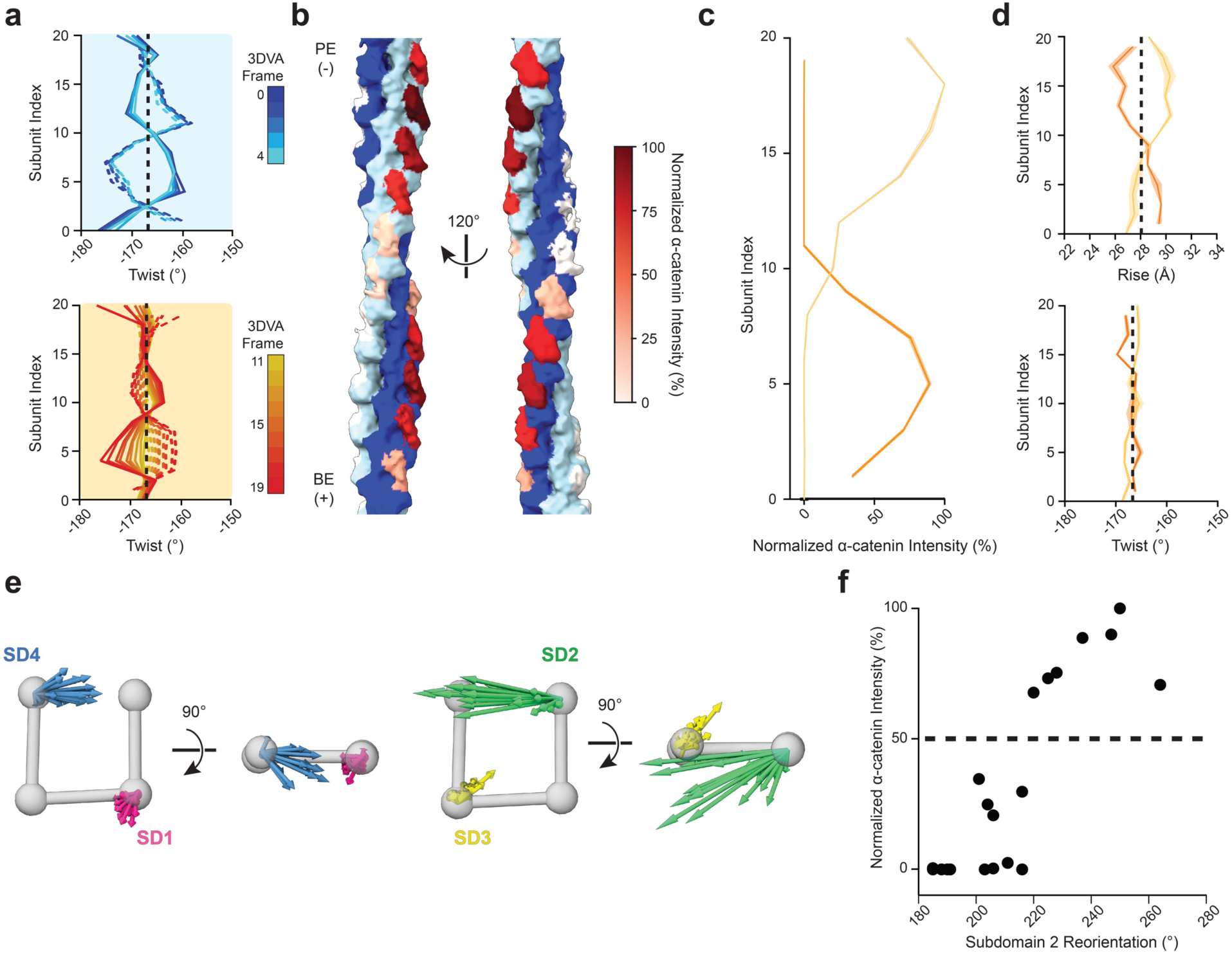
Additional analysis of the force-activated *α*-catenin–F-actin complex structure. **a**, Instantaneous helical twist of selected 3DVA frames from Fig. 4b,c. Vertical dashed lines indicate canonical F-actin twist. **b**, Orthogonal views of the post-3DVA α-catenin–F-actin complex map, colored as in Fig. 4a. **c**, Quantification of α-catenin density intensity in post-3DVA map. Yellow corresponds to light blue strand from a, and orange corresponds to dark blue strand. **d**, Instantaneous helical parameters of consensus map, colored as in b. Shaded regions represent 95% CI from 3 independent analyses. Vertical dashed lines indicate parameters of canonical F-actin. **e**, Superimposed subdomain displacement vectors versus a canonical F-actin subunit from all protomers after MDFF analysis of the consensus α-catenin–F-actin complex map, displayed as in Fig. S6. The averages of these vectors are displayed in Fig. 4f, top. **f**, Quantification of actin subdomain 2 reorientation versus α-catenin intensity. Dashed line indicates 50%.

**Table S1.** Cryo-EM data collection and processing statistics.

| Data collection |  |  |  |  |  |  |  |
| --- | --- | --- | --- | --- | --- | --- | --- |
| Microscope | Titan Krios |  |  |  |  |  |  |
| Voltage (kV) | 300 |  |  |  |  |  |  |
| Sample | Dual Motor +ATP | | | Dual Motor -ATP | | Dual Motor +ATP + α-catenin | |
| Detector | K3 | K3 | K3 | K3 |  | K2 summit |  |
| Magnification | 64,000 | 64,000 | 64,000 | 64,000 |  | 22,500 |  |
| Electron exposure ($e^-/\text{\AA}^2$) | 59.156 | 59.156 | 59.156 | 59.156 | | 56.53 | |
| Exposure rate ($e^-/\text{pixel/s}$) | 23.66 | 23.66 | 23.66 | 23.66 | | 5.65 | |
| Calibrated pixel size ($\text{\AA}$) | 1.08 | 1.08 | 1.08 | 1.08 | | 1.33 | |
| Defocus range ($\mu\text{m}$) | -1.5 to -3.5 | -1.5 to -3.5 | -1.5 to -3.5 | -1.5 to -3.5 | | -1.5 to -3.5 | |
| Micrographs (No.) | 12,453 | 10,842 | 8,504 | 3,749 |  | 4,455 |  |
| Data Processing | Dual Motor +ATP |  |  | -ATP<br>Control 1 | -ATP<br>Control 2 | Consensus | 3DVA<br>sorted |
| Initial particles (No.) | 276,493 |  |  | 345,515 |  | 133,677 |  |
| Final particles (No.) | 13,146 |  |  | 3436 | 3137 | 3,289 | 822 |
| Symmetry imposed | C1 |  |  | C1 | C1 | C1 | C1 |
| Map resolution ($\text{\AA}$) | 9.5 | | | 9.1 | 8.9 | 10.4 | 12.3 |
| FSC threshold | 0.143 |  |  | 0.143 | 0.143 | 0.143 | 0.143 |
| Map sharpening B factor ($\text{\AA}^2$) | -886.6 | | | -850.8 | -801.5 | -1151.9 | -100.0 |

**Table S2.** Coarse grained molecular dynamics simulation parameters.

|  |  |
| --- | --- |
| $k_{l\text{ axial}}$ (pN/nm) | 265 |
| $k_{l\text{ lateral}}$ (pN/nm) | 53 |
| $k_{\theta 1}$ (kJ) | $1.2 \times 10^{-15}$ |
| $k_{\theta 2}$ (kJ) | $2.4 \times 10^{-15}$ |
| $k_{\theta 3}$ (kJ) | $1.2 \times 10^{-15}$ |
| $k_{\theta 4}$ (kJ) | $6 \times 10^{-16}$ |
| $k_{\theta 5}$ (kJ) | $6 \times 10^{-16}$ |
| $k_{\theta 6}$ (kJ) | $3 \times 10^{-16}$ |
| $k_{\phi 1}$ (kJ) | $1.5 \times 10^{-16}$ |
| $k_{\phi 2}$ (kJ) | $7.5 \times 10^{-16}$ |

## Supplementary Video Legends

**Video S1: Cryo-ET of a Ptk2 cell adhesion marked by zyxin.** Segmentation is colored as in Fig. 1b.

**Video S2: Reconstituting myosin motility on cryo-EM grids.** Unanchored actin filaments were visualized by rhodamine-actin fluorescence (blue) in the presence of the indicated motors, while the holey carbon substrate was visualized by brightfield (grey).

**Video S3: Dynamics of dual motor assay on cryo-EM grids.** Actin filaments were visualized by rhodamine-actin fluorescence (blue) in the absence and presence of ATP to activate motors. Arrowhead indicates a mechanically-induced severing event. The substrate was visualized by brightfield (grey).

**Video S4: Dynamics of single motor assays on cryo-EM grids.** Actin filaments anchored through biotinylated seeds were visualized by rhodamine-actin fluorescence (blue) in the presence of either myosin-6 (“pointed end”) or myosin-5 (“barbed end”). Arrowheads indicate mechanically-induced severing events. The distribution of myosins was visualized by eGFP fluorescence (grey), which marks the substrate.

**Video S5: Cryo-ET of superhelical oscillatory domain.** Serial slices through a tomogram from the barbed end directed force condition. Scale bar, 50 nm.

**Video S6: Coarse-grained MD simulations mimicking motor dynamics elicit F-actin spirals.** Movie of the time evolution of simulations presented in Fig. 2f.

**Video S7: Reconstruction of superhelical F-actin.** Movie of stitched superhelix reconstruction (colored as in Fig. 3a), highlighting the three-dimensional corkscrewing character of the filament.

**Video S8: 3DVA captures coupling between F-actin remodeling and α-catenin engagement.** Movie of 3DVA frames, highlighting both changes in filament curvature and α-catenin binding. α-Catenin is colored by normalized intensity (a proxy for occupancy) as in Fig. 4a.

## Methods

Unless otherwise noted, chemicals and buffer components were purchased from Sigma-Aldrich.

### Cell Culture

PtK2 cells (ATCC, CCL-56) were cultured in MEM (Gibco) supplemented with 10% fetal bovine serum (FBS), 1 mM sodium pyruvate, 1X non-essential amino acids (Gibco), and 1X antibiotic-antimycotic (Gibco), at 37 °C in 5% CO_2_.

### Lentivirus production and cell line generation

For generation of a doxycycline-inducible F-tractin-mScarlet and zyxin-LDO-mNeongreen PtK2 cell line, HEK293T cells were co-transfected with lentiviral plasmids and packing plasmids (PMD2.G and PsPAX2) using Lipofectamine 2000 (Thermo Fisher Scientific). 6 hours post-transfection, the culture media was refreshed. 48 hours post-transfection, the culture media containing lentivirus was harvested and filtered. Lentivirus infection with polybrene (5 μg/mL) followed immediately after harvesting. Three types of lentivirus were added to PtK2 cells: F-tractin-mScarlet, zyxin-LDO-mNeongreen, and TetOn. 24 hours post-infection, cells that stably integrated the lentiviral plasmids were selected with puromycin and blasticidin for one week.

### Grid preparation for cellular cryo-electron tomography

C-flat 1.2/1.3 holey carbon Au 200 mesh grids (Electron Microscopy Sciences) were sputter coated with 22 nm of carbon and baked in a dry oven at 60 °C overnight. Grids were plasma cleaned and coated with 10 μg/mL fibronectin (Sigma, FC010) in DPBS for 1-3 hours at 37 °C in a tissue culture incubator. Fibronectin coated grids were washed with DPBS and stored in fresh PtK2 culture medium in a 35 mm tissue culture dish for immediate use.

Doxycycline-inducible F-tractin-mScarlet and zyxin-LDO-mNeongreen PtK2 cells were grown to 90% confluency before passaging. Cells were thoroughly trypsinized and filtered through a 0.45 μm filter to remove clumps. 30,000 cells were added to each 35 mm tissue culture dish containing the fibronectin coated grids. 24 hours later, the media was replaced with fresh PtK2 media containing 100 ng/mL of doxycycline to induce expression of F-tractin-mScarlet and zyxin-LDO-mNeongreen. The next day, grids were prepared for plunge freezing. Grids for tomograms presented in Fig. 1b and Fig. S1c were treated with 1 µg/ml Rho Activator II (Cytoskeleton, CN03) for 3.5 hours, while grids for tomogram presented in Fig. S1b were untreated. The grids were then washed with DPBS and loaded onto a Leica EM GP plunge freezer operating at 25 °C. 3 μL of DPBS was added to the front of the grid, followed by blotting from the back using a Whatman no. 5 filter paper for 7 s, then flash-frozen in liquid ethane.

### Cryo-Fluorescence Imaging

Grids were mounted on a Leica Cryo-CLEM microscope. Epifluorescence microscopy was performed using a 50X 0.9 NA air objective at -180 °C. Z stacks of images were captured at a depth of 16-bit. Image acquisition was performed using LAS X software (Leica). Images were post-processed with THUNDER (Leica) to denoise, then maximum intensity projected.

### Cellular cryo-ET data collection

Data were collected on a spherical-aberration (Cs) corrected Titan Krios TEM (Thermo Fisher Scientific) operating at 300 kV and equipped with a K3 direct electron detector (Gatan) and a BioQuantum energy filter (Gatan) using a slit width of 20 eV. Micrographs were collected at a nominal magnification of 15,000X, which corresponds to a calibrated pixel-size of 5.05 Å (2.525 Å in super resolution mode). The tilt-series were collected using the FastTomo acquisition scheme^1^ in SerialEM^2^ using a target defocus of -5 μm underfocus. Tilt angles ranged from -54° to +54° in 3° increments, grouped in a dose-symmetric manner^3^. Each tilt in the tilt-series had a total acquisition time of 2.4 s and total dose of 2.82 e^-^ / Å^2^, fractionated across 16 frames (0.18 e^-^ / Å^2^ per frame).

### Cellular cryo-ET data processing and segmentation

Individual tilts were motion-corrected and binned 2X (to a pixel size of 5.05 Å) using MotionCor2^4^, and contrast transfer function (CTF) parameters were estimated using CTFFIND4^5^. Tilt-series alignment was carried out using AreTomo2^6^, and reconstructed using IMOD^7^ with back-projection at a binning of 2 (voxel size of 10.1 Å). The reconstructed tomograms were denoised using cryoCARE^8^, then the missing wedge was predicted and restored with IsoNet^9^. The tomograms were then segmented using a U-Net convolutional neural-network with the Dragonfly software^10^. For the tomogram displayed in Fig. S1b, the plasma membrane was segmented using MemBrain v2^11^. All membrane segmentations were smoothed using mean curvature motion^12^. Segmentations were visualized using ChimeraX^13^.

### Actin purification

Actin was prepared from chicken skeletal muscle as previously described^14^. All steps were performed at 4 °C unless otherwise indicated. Briefly, 1 g of acetone powder was resuspended in 20 mL of G-Ca buffer (G buffer: 2 mM Tris-HCl pH 8, 0.5 mM dithiothreitol (DTT), 0.2 M ATP, and 0.01% NaN_3_, supplemented with 0.1 mM CaCl_2_), then mixed by inversion for 30 minutes. The mixture was ultracentrifuged in a Beckman Ti70 rotor at 42,500 rpm (79,766 g) for 30 minutes. The supernatant, containing G-actin monomers, was collected, and 50 mM KCl and 2 mM MgCl_2_ were added to induce F-actin polymerization for 1 hour. 0.8 M KCl was then added and the mixture incubated for 30 minutes to dissociate contaminants from F-actin, followed by ultracentrifugation in a Ti70 rotor at 42,500 rpm (79,766 g) for 3 hours. The pellet, containing F-actin, was collected and resuspended in 2 ml G-Ca buffer, then incubated overnight to depolymerize filaments. The mixture was then Dounce homogenized for 10-15 passes, sequentially sheared with 26G and 30G needles, then dialyzed against G-Ca buffer overnight in Spectra/Por 1 dialysis tubing (6-8 kDa MWCO). The solution was then collected, sheared again with a 30G needle, then dialyzed against G-Ca buffer for an additional day. The solution, containing dissociated G-actin monomers, was collected and ultracentrifuged in a Beckman Ti90 rotor at 70,000 rpm (187,354 g) for 3 hours. The upper two-thirds of the supernatant was then loaded on to a Cytiva HiLoad 16/600 column for size-exclusion chromatography. Purified G-actin was maintained in G-Ca buffer at 4 °C.

### Protein expression and purification

Plasmids were propagated in NEB 5-alpha competent *E. Coli* cells and purified with either Qiaprep spin miniprep kits (Qiagen, for bacterial expression) or the PureYield plasmid maxiprep system (Promega, for transfection). Full-length human calmodulin was expressed in Rosetta2(DE3) *E. coli* cells and purified at 4 °C according to a published protocol^15^. The purified protein was collected in Storage Buffer (20 mM Tris-HCl pH 8.0, 100 mM NaCl, 2 mM β-mercaptoethanol, and 5% v/v glycerol), flash frozen in liquid nitrogen, and stored at -80 °C prior to use. For experiments presented in Fig. 1, Fig. S2, and Videos S2-4, constructs for murine myosin-5a (AA 1-1090) and human myosin-6 (AA 1-1021) featuring C-terminal GFP and Flag tags were expressed and purified from Sf9 insect cells using the baculovirus system as previously described^16^. In the myosin-6 construct, a GCN4 leucine zipper dimerization domain replaced the smooth muscle coiled-coil region originally employed by Nishikawa et al^17^. For all other experiments, equivalent constructs for human myosin-5a (AA 1-1090) and myosin-6 (AA 1-1021 were expressed from a modified pCAG vector featuring a C-terminal GFP and Flag tag in FreeStyle 293-F cells (ThermoFisher) and purified as described^18^.

Briefly, cells were cultured in FreeStyle expression medium (ThermoFisher) at 37 °C on an orbital shaker in the presence of 8% CO_2_. Cells were transfected when the culture reached a density of 1.8 x 10^6^ cells / mL. Per 400 mL of culture, 1.2 mL of 1 mg/mL PEI MAX (PolySciences) was premixed with 400 µg of plasmid in 15 mL of FreeStyle expression medium and incubated for 20 minutes at room temperature prior to transfection. Myosin-5a / myosin-6 were co-transfected with a modified pCAG vector containing untagged, full-length human calmodulin (AA 1-149) at a mass ratio of 1:6. Cells were cultured for an additional 60 hours, then harvested and snap frozen in liquid nitrogen. Cell pellets were stored at -80 °C.

For myosin purification, all steps were performed at 4 °C. Cells were resuspended in Myosin Lysis Buffer (50 mM Tris-HCl pH 8.0, 150 mM NaCl, 2 mM MgCl_2_, 0.2% 3-((3-cholamidopropyl) dimethylammonio)-1-propanesulfonate (CHAPS), 2 mM ATP, 1 mM phenylmethylsulfonyl fluoride (PMSF), 1 μg/mL aprotinin, leupeptin, and pepstatin) and incubated with rocking for 40 minutes. The lysate was clarified by centrifugation in a Beckman JA-25.50 rotor at 20,000 g for 30 minutes. The supernatant was collected and incubated with anti-Flag M2 affinity beads (Sigma-Aldrich) on a rocker for 1.5 hours. The beads were collected and washed three times with Myosin Purification Buffer (50 mM Tris-HCl pH 8.0, 150 mM NaCl, 2 mM MgCl_2_, and 2 mM ATP), then eluted with Myosin Purification Buffer supplemented with 100 µg/ml Flag peptide (Sigma-Aldrich). The eluted protein was buffer exchanged into Myosin Storage Buffer (10 mM Tris-HCl pH 8.0, 100 mM NaCl, 2 mM MgCl_2_, and 3 mM DTT) using an Amicon Ultra-4 concentrator (50 kDa MWCO), then snap frozen in liquid nitrogen. Purified myosins were stored at -80 °C prior to use.

The C-terminal actin-binding domain (ABD) of human α-catenin (AA 664-906) was expressed from a pET vector encoded an N-terminal 6XHis tag, strep tag, and TEV protease cleavage site as previously described^19^. Transformed Rosetta2(DE3) *E. Coli* were cultured in LB media at 37 °C to an optical density of 0.8-1.0, then induced with 0.7 mM isopropyl β-D-1-thiogalactopyranoside (IPTG) and cultured at 16 °C for 16 hours. Cells were then harvested and flash frozen in liquid nitrogen. Cell pellets were stored at -80 °C.

For purification, all steps were performed at 4 °C. Cells were resuspend in Lysis Buffer (50 mM Tris-HCl pH 8.0, 150 mM NaCl, 2 mM β-mercaptoethanol, 20 mM imidazole) and disrupted in an Avestin Emulsiflex C5 homogenizer. The lysate was clarified at 15,000 g in a Beckman JA-25.50 rotor for 30 minutes, then the supernatant was collected and incubated with Ni-NTA resin (Qiagen) with rocking for 1 hour. The resin was collected, washed with 5 bed volumes of Lysis Buffer, then eluted with Elution Buffer (50 mM Tris-HCl pH 8.0, 150 mM NaCl, 2 mM β-mercaptoethanol, 300 mM imidazole). Purified His-tagged TEV protease (prepared according to a published protocol^20^) was then added to reach a 0.05 mg/ml working concentration, and the solution was dialyzed against Dialysis Buffer (20 mM Tris-HCl pH 8.0, 300 mM NaCl, 2 mM β-mercaptoethanol) for 16 hours. The sample was then collected and re-applied to Ni-NTA resin to remove TEV protease. The flowthrough was collected, then sequentially purified by anion exchange chromatography using a HiTrapQ HP column (Cytiva), followed by size exclusion chromatography on an Superdex 200 Increase column (Cytiva) in Gel Filtration Buffer (20 mM Tris-HCl pH 8.0, 100 mM NaCl, 2 mM β-mercaptoethanol). The protein was snap frozen in liquid nitrogen and stored at -80°C until use.

### Reconstituting myosin force generation on cryo-EM substrates

All steps were performed at room temperature unless otherwise noted. Filamentous (F)-actin was prepared from G-actin monomers in G-Mg (G Buffer supplemented with 0.1 mM MgCl_2_) plus KMEI buffer (50 mM KCl, 1 mM MgCl_2_, 1 mM ethylene glycol-bis(β-aminoethyl ether)-N,N,N′,N′-tetraacetic acid (EGTA), 10 mM imidazole pH 7.0, 1 mM DTT) as previously described^21^. CF-1.2/1.3-3Au 200-mesh gold C-flat holey carbon cryo-TEM grids (EMS, CF213-50-Au) were used for initial cryo-EM imaging and tomography studies, while CF-1.2/1.3-3Au 300-mesh grids (EMS, CF313-50-Au) were used for single-particle studies.

For the dual motor condition, the following solutions were prepared at 4°C in Motility Buffer (MB, 20 mM 3-(*N*-morpholino)propanesulfonic acid (MOPS) pH 7.4, 5 mM MgCl_2_, 0.1 mM EGTA, 1 mM DTT): MB-PVP, 0.1% polyvinylpyrrolidone (Sigma-Aldrich, PVP10); MB-anchor, 0.1 mg/mL mouse monoclonal anti-GFP antibody (Sigma-Aldrich, G6539); MB-myosin, 0.02 μM myosin-5 and 0.08 μM myosin-6; MB-ATP, 1 µM ATP and 0.01% Nonidet P-40 Substitute (NP-40, Roche); MB-no-ATP, 0.01% NP-40. Solutions were clarified by ultracentrifugation at 50,000 rpm (108,726 g) in a Beckman TLA-100 rotor, then brought to room temperature immediately prior to grid preparation.

Grids were untreated prior to sample preparation, as we found plasma cleaning and other treatments reduced the activity of myosins. First, 6 μL of MB-anchor was applied to each side of the grid and incubated for 2 minutes. 6 μL of MB-myosin was then applied to each side of the grid and incubated for an additional 2 minutes. 20 μL of MB-PVP was then applied to the top (shiny) side of the grid and incubated to block the surface for 1 minute. 6 μL of F-actin was then applied and incubated for 40 s. The grid was then washed in a 500 μL reservoir of MB in a microcentrifuge tube, then placed in the chamber of a Leica EM-GP plunge freezing device operating at 25 °C and 100% humidity. 6 μL of either MB-ATP (+force generation) or MB-no-ATP (no force generation) was then applied and incubated for 40 s. The grid was then blotted for 5 s and plunged into liquid ethane. For each grid preparation session, samples with F-actin concentrations ranging between 0.4-1.0 μM were prepared and screened, and specimens featuring optimal filament density were selected for further analysis.

For the barbed-end directed and pointed-end directed single motor conditions, reagents were prepared as above, with the following modifications: MB-anchor was supplemented with 500 nM streptavidin (VWR, S000-01), and two separate myosin buffers were prepared, MB-myosin-5 (0.02 μM myosin-5 in MB) and MB-myosin-6 (0.08 μM myosin-6 in MB). To prepare 25% biotin-F-actin seeds biotinylated actin monomers (Cytoskeleton, AB07-C) were co-polymerized at a 1:4 molar ration with unlabeled monomers with a total concentration of 10 μM for 1 hour, then sheared through a 30G needle. F-actin was then extended by mixing seeds with unlabeled monomers at a 1:5 molar ratio and polymerizing for 1 additional hour. Samples were then prepared as above, using either MB-myosin-5 (barbed-end directed) or MB-myosin-6 (minus-end directed).

Samples for the force-activated α-catenin–F-actin complex were prepared identically as for the dual motor condition, with the following modifications. After mounting in the plunge-freezing apparatus, 3 μL of MB-ATP supplemented with α-catenin ABD was applied and incubated for 30 s, then the grid was blotted for 5 s and plunged into liquid ethane. Grids were prepared with varying F-actin / α-catenin ABD concentrations, and 0.6 μM F-actin / 1 μM α-catenin ABD was found to be optimal after screening. All grids were stored in liquid nitrogen prior to cryo-EM data collection.

### Fluorescence imaging of myosin activity on cryo-EM substrates

For data presented in Videos S2-4, samples were prepared as described above, except 10% rhodamine G-actin (Cytoskeleton, AR05-B) was included in all polymerization reactions (F-actin and biotinylated seeds). Instead of plunge-freezing, grids were sandwiched between two No. 1.5 24 x 60 mm glass coverslips (Corning) in a drop of MB-ATP supplemented with oxygen scavengers (20 μg/mL catalase, 100 μg/mL glucose oxidase, 125 mg/mL glucose, 50 mM DTT). Epifluorescence image sequences (movies) were collected at room temperature using a 60X 1.40 NA PlanApo oil-immersion objective lens (Nikon) on an inverted TE2000-E microscope (Nikon) equipped with a solid-state white light illumination system (Lumencor SOLA SE). Image sequences (movies) were collected on a Photometrics Coolsnap HQ2 CCD camera with a frame rate of one exposure every 2 or 3 s, using filters for visualizing rhodamine (excitation 650 / 25, emission 607 / 36). A single image (either brightfield to directly visualize the cryo-EM grid, or fluorescence using filters to visualize anchored, GFP-tagged myosins: excitation 485 / 20, emission 525 / 30) was taken before and after each image sequence to visualize the grid substrate and verify the lack of substantial drift.

### Low magnification cryo-EM of oscillatory domains

For data and analysis presented in Fig. 1d-f and Fig. S2a-c, samples were imaged on a Tecnai F20 transmission electron microscope operating at 120 kV using a Gatan 626 cryo-transfer holder, operated with the Leginon software package^22^. Exposures were collected at 29,000X magnification (corresponding to a calibrated pixel size of 3.2 Å / pixel) on a Gatan Ultrascan 4000 CCD camera with –4 µm underfocus. Images were low-pass filtered to 20 Å and binned by 4 for visualization / analysis. Oscillatory domain wavelengths were manually quantified with FIJI^23^.

### Cryo-electron tomography of reconstituted specimens

Grids were clipped and transferred to Talos Arctica TEM (ThermoFisher) operating at 200 kV with a Gatan K2 Summit camera. Tomograms were collected using a dose symmetric scheme from –60° to +60° degrees with 3° angular increments. Each tilt image was fractionated into 10 frames with a total exposure dose of 2.4 e^-^ / Å ^2^ (0.24 e^-^ / Å ^2^ / frame) and a total exposure time of 1 s. The defocus range was between -1.5 μm and -5 μm underfocus. The acquisition magnification was 17,000X with a pixel size of 2.4 Å. Frames were aligned with MotionCor2^4^.

Tomograms were reconstructed using the Appion-Protomo software^24,25^. Images were aligned and dose weighted, then reconstructed using the Simultaneous Iterative Reconstruction Technique (SIRT) as implemented in Tomo3D^26^. Reconstructions were binned by 4 resulting in a final pixel size of 9.6 Å.

### Synthetic data generation for *in vitro* tomogram denoising

*In vitro* tomograms were processed using a denoising autoencoder (DAE) approach to enable direct, confident tracing of F-actin filaments in 3D. A DAE with a similar architecture to that reported in our previous work^27^ was trained. Briefly, synthetic datasets approximating subtomograms were first generated. 24,000 synthetic noiseless subtomograms were prepared which contained between zero and five *in silico* actin filament models (PDB: 7R8V) of varying curvature. All of these filament models were bent in a single plane (to prevent superhelix model bias), then converted to volumetric data using the pdb2mrc procedure in EMAN2^28^. Each actin filament was rotated about the rot and psi angles by random, uniformly sampled values between 0° and 359°, while the tilt rotation angle was sampled from a Gaussian probability distribution centered at 90° with a standard deviation of 20°. Each actin filament was translated in the box along each dimension by random values uniformly sampled within the range of ± 196 Å.

Matched synthetic noisy volumes were generated by first projecting the sum of the oriented filaments at integer angular increments (held constant for a particular subtomogram) uniformly sampled between 1° and 5°, inclusive. The maximum tilt was sampled from a random integer between 45° and 60°, yielding tilt angle ranges from ±45° to ±60°. These projections were corrupted by a CTF, using parameters matching those of the microscope and a defocus range of -3 μm to -11 μm underfocus, then reconstructed back into a three-dimensional volume using reconstructor class functions as implemented in the EMAN2 python package^28^. Empirical noise was modeled by extracting 1,000 subtomograms of 96 x 96 x 96 voxels from each experimental tomogram and computing the per tomogram average Fourier transform. For each synthetic volume, one of these empirical noise boxes was randomly selected, multiplied element-wise by a white noise box of the same dimensions, normalized, and scaled by a random scale factor that modulated the signal-to-noise ratio of each synthetic subtomogram. The synthetic volume was then summed with its noise volume in Fourier space. To account for interpolation artifacts from CTF application or noise addition in Fourier space, the synthetic volumes were then cropped to 48 voxels in real space. After these operations were complete, each noisy particle was paired with its corresponding noiseless ground truth particle.

### Neural network training and *in vitro* tomogram denoising

A denoising auto-encoder (DAE) featuring adversarial training was constructed and trained using an approach we have recently reported^27^. Pre-training of this denoising autoencoder consisted of 3D convolutional layers in a U-net architecture, and it was performed using a single NVIDIA A100 GPU with 80 GB of VRAM, using a learning rate of 0.00001. Training was run on the 24,000 pairs of synthetic noisy and noiseless volumes, with a 90 : 10 training : validation split, for 30 epochs. This model had a cross-correlation coefficient validation loss of 0.9147. After pre-training, an additional network head was added for domain classification by forking the network output after the feature extraction layers. The domain classification head consists of a gradient reversal layer and additional 3D convolutional layers, which are followed by flattening to a dense layer, then a binary classification layer with a sigmoid activation function. 1,000 synthetic volumes and 1,000 volumes extracted from the experimental tomograms were used as the training set for the domain classifier. Adversarial training was performed by alternatively passing these data through the domain classifier head, followed by re-training the feature extractors with the denoising head using only the 1,000 synthetic volumes. This adversarial training was run for ten iterations, and the final iteration of adversarial training was used for denoising.

The adversarial-trained neural network was then used to denoise empirical tomograms, using A100 GPUs. 48-voxel tiles sampled every 16 voxels in each dimension were extracted, normalized, and passed as inputs to the neural network. The network outputs were stitched together via maximum intensity projection.

### Quantification of oscillatory domains from *in vitro* tomograms

Denoising the *in vitro* tomograms enabled 3D contour tracing of F-actin. Denoised tomograms were visualized in UCSF Chimera^29^, and each filament’s central axis was manually traced using Chimera’s marker set tool. Univariate splines were fit through the data with a fixed smoothing factor of 150 to minimize jitter from manual picking. The spline fits were resampled evenly with 9.6 Å steps and visually inspected to ensure trace quality.

The cartesian coordinates of these traces were subsequently analyzed. The instantaneous curvature and torsion along each filament trace was inspected to identify filament segments with potential oscillatory character. Filament segments with high, oscillating curvature and torsion that were between 350 nm and 500 nm were identified and selected. Principal component analysis (PCA) was performed on these selected filament segments, and the decompositions were aligned such that the maximum amplitude along the second principal component was at the center point and the second derivative of the second principal component at this point was positive. If a filament’s principal component decomposition was inverted from the other traces, the starting end of the filament spline was flipped. One limitation of this analysis is that filament polarity could not be assigned in the denoised tomograms. The aligned traces were analyzed within a common 350 nm window to produce average traces, as well as amplitude and wavelength measurements (Fig. 2c,d).

To assess whether the superhelical filament segments exhibited nonuniform orientation relative to the ice layer, the second principal component’s angle relative to the ice layer was analyzed. The tomograms contained protein aggregates at the air-water interface, which appeared as amorphous density in the denoised tomograms. Chimera’s marker set tool was used to position centroids within these densities. PCA was then performed on these points, where the first and second principal components defined the ice plane. The angle between the second principal component of the oscillatory region (i.e. the major axis of each superhelix’s elliptical cross-section) and the ice plane was measured by projecting the ice plane’s normal vector along the first principal component of the oscillatory domain and computing the angle between these vectors. A Kolmogorov–Smirnov test as implemented in Scipy^30^ was performed to assess whether the observed major axis angle distribution was significantly different from a normal distribution, and no significant difference was found (p = 0.27).

### Coarse-grained molecular dynamics simulations

Coarse-grained molecular dynamics simulations of individual actin filaments under force were performed using the software package ESPResSO^31^ and custom Python scripts. An actin filament composed of 400 subunits was modeled as a 3D network of springs, with each subunit being approximated by a spherical particle that connects to its neighbors via springs (Fig. 2e). The energetics of the actin filament is described by the harmonic bond potential (Equation 1), harmonic angle potential (Equation 2), and dihedral potential (Equation 3):

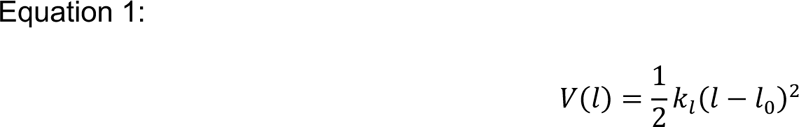

Where *k_l_* is the harmonic bond stiffness, and *l*_0_ is the lateral or axial bond length in equilibrium configuration.

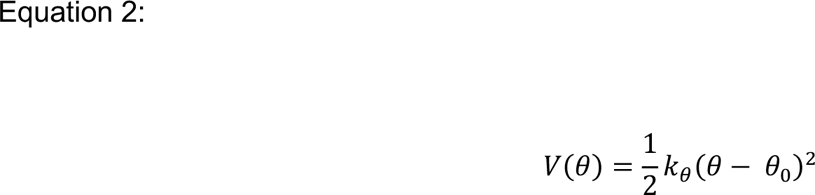

Where *k_θ_* is the harmonic angle bending constant, and *θ*_0_ is the angle formed by neighboring triplets in equilibrium configuration. Each particle is associated with six different harmonic angles.

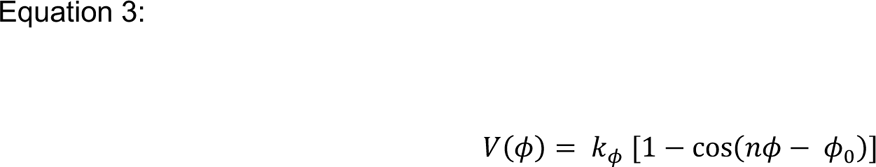

Where *k_ϕ_* is the dihedral angle bending constant, *ϕ*_0_ is the dihedral angle formed by neighboring quadruplets in equilibrium configuration, and *n* is the multiplicity of the potential (number of minima). Each particle is associated with two different dihedral angles; *n* was set to 1 for simplicity.

The geometric parameters of equilibrium configuration, *l*_0_, *θ*_0_, and *ϕ*_0_ were derived from the canonical actin filament structure^32^, which depicts two right-handed helices wrapping around each other, leading to a left-handed helix with a helical rise of 27.8 Å and twist of -166.67°. For simplicity, deformations of individual protomers were ignored. The effect of actin nucleotide state transitions on filament mechanics was also neglected.

The stiffness and bending constants, *k_l_*, *k_θ_*, and *k_ϕ_*, have not been experimentally measured; nonetheless, they can be calibrated through matching to previously reported data, as has been done by Yogurtcu et al.^33^. Chu and Voth have established a force-extension relationship via coarse-grained molecular dynamics simulation of the actin filament, revealing a stretch stiffness of 37 pN / nm and 31 pN / nm for actin filaments in the ATP and ADP state, respectively^34^. The extensibility of an actin filament is determined by the harmonic bond potential and harmonic angle potential.Therefore, to calibrate *k_l_* and *k_l_*, a parameter scan was performed by simulating an actin filament composed of 39 subunits (the same number of subunits modelled by Chu and Voth^35^) under tension in vacuum, followed by extracting the stretch stiffness by plotting the pulling force against the filament extension at equilibrium (Fig. S2f).

The capacity of an actin filament to over- and under-twist is mainly determined by the dihedral potential, which is modulated by the harmonic angle potential due to twist-bend coupling^36^. Recently, Bibeau et al. reported the cumulative variance of twist as a function of subunit index observed in cryo-EM data^37^. To calibrate *k_ϕ_* and further refine *k_l_*, an additional parameter scan was performed by simulating the thermal fluctuations of an actin filament composed of 100 subunits using the Langevin thermostat with the friction coefficient of water at room temperature, followed by establishing the subunit index dependence of cumulative twist variance (Fig. S2g). To calculate subunit twist, the centroids of consecutive particles on the left-handed helix were found, neighboring centroids were connected by line segments, then the midpoints of neighboring line segments were connected by a final set of line segments. Each pair of consecutive subunits was projected to the nearest final line segment. The twist was then calculated from the projection vectors. For each timepoint, the twist deviation from canonical F-actin for each subunit was calculated. Then, a zero-mean normal distribution was fitted to the distribution of these twist deviations for each subunit across the simulation. The variance for each subunit was derived from the fit and plotted against subunit index (Fig. S2g). The initial and terminal 20 subunits were excluded from the analysis to avoid edge effects. The final parameters are reported in Table S2.

The dynamics of an actin filament under force were simulated by fixing the initial five subunits and applying pulling or pushing force on the terminal five subunits (across both strands). The magnitude of force applied to each particle was 5 pN, approximating the force exerted by a single myosin and adding up to a total force of 25 pN. To define the central axis, the same process for calculating subunit twist was implemented, followed by fitting a 3D spline along the final set of midpoints. Principal component analyses were performed on the 3D spline fits (Fig. 2f). The period of the spline fits was extracted from the position of the first peak in the autocorrelation function along PC1, computed with a Gaussian blur (σ = 2 nm; Fig. 2g). Spline fits with an amplitude of less than 15 nm were considered insignificant and thus were excluded from the period analyses.

### Single particle cryo-EM data acquisition

For the dual motor condition cryo-EM data were recorded on a Titan Krios (ThermoFisher/FEI) operated at 300 kV equipped with a Gatan K3 camera, BioQuantum energy filter and spherical aberration (Cs) corrector. SerialEM^2^ was used for automated data collection. Movies were collected at a magnification of 64,000X in super-resolution mode resulting in a calibrated pixel size of 1.08 Å / pixel (superresolution pixel size of 0.54 Å / pixel), over a defocus range of -1.5 to -3.5 µm underfocus. 63 frames were recorded over 2.5 s of exposure at a dose rate of 27.6 e^-^ / pixel / s (23.7 e^-^ / Å^2^ / s) for a cumulative dose of 59.2 e^-^ / Å^2^. Three datasets were collected with these imaging conditions consisting of 12,453 movies, 10,842 movies, and 8,504 movies, respectively. The first dataset was used for preliminary processing, and all three datasets were pooled to generate the final reconstruction. A single -ATP control dataset was collected under the same imaging conditions, consisting of 3,749 movies.

For the force-activated α-catenin–F-actin complex, cryo-EM data were recorded on a Titan Krios (ThermoFisher/FEI) operated at 300 kV equipped with a Gatan K2 Summit camera. SerialEM^2^ was used for automated data collection. Movies were collected at a nominal magnification of 22,500X in super-resolution mode, resulting in a calibrated pixel size of 1.33 Å / pixel (super-resolution pixel size of 0.665 Å / pixel), over a defocus range of -1.5 to -3.5 µm underfocus. 50 frames were recorded over 10 s of exposure at a dose rate of 10 e^-^ / pixel / s (e^-^ / Å^2^ / s) for a cumulative dose of 56.5 e^-^ / Å^2^. Three datasets were collected with these imaging conditions consisting of 1,909 movies, 1,792 movies, and 754 movies, respectively. The datasets were pooled prior to processing.

### Micrograph pre-processing

Movies were aligned with MotionCor2 using 5 × 5 patches^4^, and dose-weighting sums^38^ were generated from twofold binned frames with Fourier cropping, resulting in a pixel size of 1.08 Å in the images collected in the dual motor condition datasets (+/- ATP) and 1.33 Å for the force-activated α- catenin–F-actin complex datasets. Non-dose-weighted sums were used for CTF parameter estimation using CTFFIND4^5^.

### Neural network architecture and training for cryo-EM data processing

To analyze single-particle data, a DAE featuring a previously described architecture^32^ was trained to learn features of actin filaments in cryo-EM projection images (Fig. S3a). In brief, the DAE consisted of an encoder and decoder. The encoder was composed of nine convolutional layers followed by three dense, fully-connected layers of decreasing size. The decoder was composed of two dense, fully-connected layers of increasing size, which then connected to nine convolutional layers, the last of which produced the denoised image.

Synthetic projection images to train the DAE and an accompanying semantic segmentation network were generated as outlined previously^32^. To improve network performance for semantic segmentation of the experimental data presented in this study, two modifications were made: incorporation of known background picks, and improvement of the noise model. Our data collection scheme resulted in a substantial amount of carbon in the micrographs. To prevent the semantic segmentation network from picking the edges of holes or actin filaments over carbon, 15,208 manual picks of hole edges and thick carbon areas were selected from micrographs in the dataset using RELION^39^. These picks were extracted at a box size of 512 pixels, then binned by 4 to a box size of 128 pixels and a pixel size of 4.32 Å / pixel. These picks were then integrated into the training dataset of the fully convolutional neural network for semantic segmentation, paired with targets composed entirely of background with no signal.

In our previously reported neural network training schemes, EMAN2’s pink noise generation function^28^ was used to produce realistic-looking projection images. However, this noise model proved insufficient to pick on the current datasets, possibly due to ice thickness or different microscope / detector parameters. To improve the approximation of the synthetic data to real micrographs, an empirical pink noise model was used. The synthetic 2D particle image generation model can be summarized by equation 4:

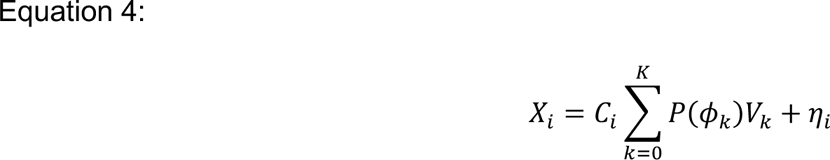

The following procedure was used to generate a particle image *X_i_*: first, a synthetic volume was randomly chosen from a library of 135 bent actin filaments *V_k_*, and the pose of the filament was transformed by a 3D rotation and translation before it was projected *P*(*ϕ_k_*). This process was repeated for each filament in the image (*K*, up to four filaments per particle image), and the results were summed in real space to produce a noiseless projection image. Then, the image was corrupted by a CTF with the sampled defocus range, C_s_, amplitude contrast, and voltage matching those used for CTF estimation. Finally, noise was added in Fourier space *η_i_*.

The noise model, *η*, was empirically determined by using RELION to manually pick and extract 2,177 empty regions within holes in the micrographs to yield empty noise boxes with pixel sizes of 4.32 Å per pixel and box sizes of 128 pixels. The average and standard deviation of the Fourier transform of each empty pick was computed to model the empirical noise distribution. For each synthetic particle image, a white noise box was generated, and the white noise box’s Fourier transform was multiplied element-wise by the sum of the empirical noise average and a Gaussian sampling of the empirical noise standard deviation. This pink noise box, *η_i_*, was added to the Fourier transform of the image after CTF corruption, and the inverse Fourier transform was computed to yield the final noisy projection image. To produce the semantic segmentation target, the noiseless projection image was lowpass filtered to 40 Å, binarized, and eroded by 8 pixels.

To train the DAE, 800,000 image pairs with a 90:10 training/validation split and a learning rate of 0.00005 were used to train the network. Cross correlation coefficient was used as the loss function. The network was trained until validation loss did not improve for three epochs; then the best network weights were restored and saved. The validation CCC of the trained DAE was 0.9887.

The trained convolutional layers of the DAE were then used to initialize the weights of the initial layers of a fully convolutional neural network for semantic segmentation, featuring a previously described architecture^32^. Briefly, the trained convolutional encoding layers of the DAE were copied as a separate neural network, and additional convolutional layers were added to form a fully convolutional neural network for semantic segmentation^40^. The final layer of this network consisted of two channels with sigmoid activation and default initializations. To train the network for semantic segmentation, 75,000 image pairs (60,000 of which contained synthetic projections and 15,000 of which were carbon picks, as described below) with a 90:10 training/validation split and a learning rate of 0.001 were used to train the network. Binary cross-entropy (BCE) was used as the loss function. The network was trained until validation loss did not improve for three epochs; then the best network weights were restored and saved. The validation BCE of the trained DAE was 0.0444. The architectures described above have since been superseded by a U-net architecture, which we found produces better segmentation loss with a smaller training set in a shorter time^41^.

### Neural network particle picker

A custom Python script was used to pass images to a fully convolutional neural network for semantic segmentation and execute curvature-sensitive filament picking, modified from our previously described method^32^. Briefly, each micrograph was binned by 4 to a pixel size of 4.32 Å per pixel, then 128-pixel tiles featuring 32 pixels of overlap were extracted and passed as inputs to the network. The outputs were stitched together by maximum intensity projection at the overlaps, producing a semantic segmentation map of the micrograph. These maps were then binarized using a fixed, empirically determined threshold of 0.3 and skeletonized. An additional step of dilation by 8 pixels after the skeletonization was performed to link filament ends, which we found were often disjointed in these datasets. Another round of skeletonization was then performed. Branches shorter than 8 pixels were pruned, and pixels within a radius of 16 pixels from filament intersections were removed. Continuous filaments were then identified by matching tracks with common end points, and 2D splines were fit through the filaments for curvature estimation. To prevent spuriously high curvature values due to edge effects, the terminal 50 pixels of the spline were omitted from picking. From the remaining filament sections, the instantaneous curvature was measured along the spline at three-pixel intervals and used for segment selection. This resulted in substantial filament segment overlap during initial picking; these overlapping segments were retained for initial alignment before subsequent duplicate removal. Segments from the same filament were flagged in the output metadata (a RELION-formatted STAR file).

To be assigned as a superhelical segment, substantial curvature in both positive and negative directions along the same filament were required. An identified filament was considered superhelical if it contained a segment at least 200 Å in length (approximately the length of 7 subunit rises) where the instantaneous curvature was always greater than or equal to 1.5 μm^−1^, as well as another segment of the same length where the instantaneous curvature was always less than or equal to -1.5 μm^−1^ (Fig. S3b).

These criteria exclude straight filaments and filaments with substantial uniplanar curving in one direction. All filament segments assigned as superhelical were retained for subsequent image processing.

### Modeling of curvature distributions

The curvature distribution of free, unloaded F-actin would be expected to follow a Boltzmann distribution set by the F-actin’s persistence length and the filament segment length, as we modeled in our previous study^32^. In this scheme, the energy to bend F-actin is defined as:

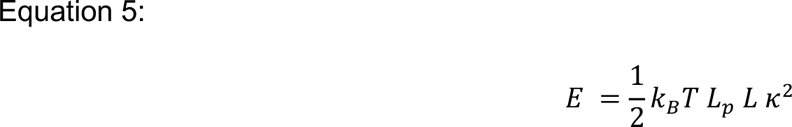

Where is the *k_B_* is the Boltzmann constant, *T* is absolute temperature, *L_p_* is persistence length in microns, *L* is the filament segment length in microns, and *κ* is curvature in inverse-microns^32,36^. This corresponds to the probability function defined by:

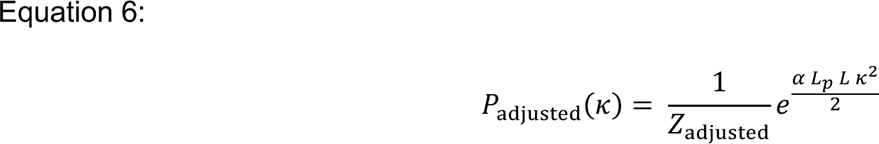

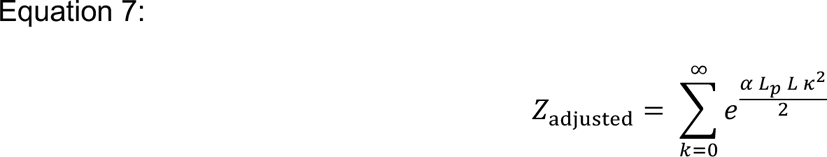

Where *α* is a multiplicative adjustment factor that serves as a proxy for differing persistence length in this simple approximation. *α* was both fixed at a value of one for a basic modeling (Fig. S3c, grey curves) and fit to the experimental curvature distributions (Fig. S3c, pink and blue curves).

### Superhelical F-actin reconstruction

For initial model generation, data from a single imaging session were used. Selected superhelical segments were extracted in RELION (v3.1.2) with a box size of 512 × 512 pixels and pixel size of 2.16 Å per pixel (bin 2; Fig. S4). To avoid reference bias, these segments were imported to cryoSPARC^42^ (v.3.2.0) for successive rounds of *ab initio* initial model generation and subsequent three-dimensional classification. The initial 148k picked segments were used for *ab initio* model generation with three classes as implemented in cryoSPARC. The subset of filament segments contributing to the best class were then selected and used for three more rounds of *ab initio* model generation with two classes; after each round only the segments belonging to the best class were retained. The reconstructions from the final round of *ab initio* model generation were then used as references for a two-class heterogeneous refinement in cryoSPARC, with the initial, unaligned 148k picked segments used as inputs. This heterogeneous refinement produced one class with clear protomer definition and one class with apparently mixed filament polarity. The segments from the good class were retained for subsequent homogeneous refinement in cryoSPARC and 3D auto-refinement in RELION. Focused 3D classification without segment alignment was then performed using three classes. Segments from a single class with poor protomer definition were rejected, and the remaining segments were refined using local 3D auto-refinement in RELION to produce the initial model for processing the full dataset.

For processing of the full dataset of 276k filament segments from three datasets, selected superhelical segments were extracted as described above. These segments were imported to cryoSPARC for 2D classification. After removing junk classes and exceptionally straight filament classes, 173k segments remained. These segments were passed to a cryoSPARC homogeneous refinement for initial alignment. The previously obtained 3D reconstruction was lowpass-filtered to 30 Å and used as the initial model. After exporting a RELION STAR file containing the cryoSPARC alignment parameters, the original filament tube IDs were restored using a custom python script. A second custom python script implementing per-filament non-maximum suppression was used to remove overlapping segments from the same filament that were closer than 768 pixels at bin 1 (approximately 830 Å), yielding 13,146 filament segments. These segments were re-imported to cryoSPARC and a nonuniform refinement^43^ was performed to improve local resolution near the filament edges using a mask encompassing the filament segment without overlap (75% z-length). Three-dimensional variability analysis (3DVA^44^) was also performed in cryoSPARC using default parameters except for the filter resolution, which was set to 8 Å.

Two independent asymmetric reconstructions from the -ATP control dataset were generated using a similar approach (Fig. S5). To obtain these reconstructions, 345k segments from the -ATP dataset that did not meet the superhelix criteria were extracted in RELION (v3.1.2) with a box size of 256 x 256 pixels and pixel size of 4.32 Å per pixel (bin 4). These segments were imported to cryoSPARC for 2D classification. After removing junk, 315k segments remained, and they were then split into two equally-sized subsets. Each subset was then subjected to the following processing steps. First, *ab initio* reconstruction of one class was performed, followed by homogeneous refinement. The aligned segments were then re-extracted in RELION at a box and pixel size matching that of the superhelix reconstruction. Overlapping segments within 830 Å were then removed, resulting in 3,436 segments in the first subset and 3,137 segments in the second subset. 3D auto-refinement with local angular searches was performed in RELION. These aligned segments were then imported to cryoSPARC for a final non-uniform refinement using a 75% z-length soft mask.

### Force-activated α-catenin–F-actin complex reconstruction

For the three pooled α-catenin datasets, particle picking was largely similar to the superhelix dataset (Fig. S7). Briefly, the micrographs were binned to a pixel size of 4.32 Å / pixel (bin 3.25) and picked using the same selection criteria to identify filaments that contained long stretches of high curvature in both directions. 133,677 segments with no overlap within 3 Å were picked, then extracted in RELION with a box size of 384 x 384 pixels and pixel size of 2.66 Å (bin 2). These extracted segments were subjected to initial 2D classification in RELION to remove junk picks. In order to align the psi (in-plane rotation) angle to a common reference, the remaining 110k segments were subjected to 3D classification in RELION using the initial superhelix model lowpass-filtered to 30 Å as a reference and an angular sampling interval of 3.7 degrees. The resulting reconstruction deviated substantially from the reference map. Therefore, the Euler angles were reset such that the rot and tilt angles were removed, the tiltPrior was set to 90°, and the psi angle was set as a prior, while translations were retained. 3D classification was performed with global rot search, local tilt and psi searches with a 12° angular search range, and bimodal priors on the psi angle.

After segments were aligned into this one class, subsequent 3D classification was performed with three classes using a fine 1.8° angular sampling interval, a global search for rot, and a 9° angular search range for tilt and psi. This yielded one bare class, which was rejected from further processing, as well as two classes with α-catenin decorating one side of the filament. To retain the maximum number of segments, the aligned segments from both decorated classes were subjected to focused 3D classification with no alignment, using a mask for α-catenin binding sites along one side of the filament. This resulted in one decorated class and one undecorated class. The decorated class was passed to a 3D auto-refinement job, and the undecorated class was subjected to another round of focused 3D classification with no alignment using a mask for α-catenin binding sites along the other side of the filament. This resulted in one undecorated class and one decorated class. The decorated class was retained and subjected to 3D auto-refinement. Upon inspection, the two refined maps were indistinguishable after a shift and rotation of one short-pitch helical step. The constituent segments of one of the classes were therefore rotated and translated by one short-pitch helical step, then combined with the segments from the other class. Overlapping segments within 768 Å were removed and refined in cryoSPARC using non-uniform refinement.

The aligned segments from the refinement were then subjected to 3DVA as implemented in cryoSPARC^44^. In order to detect large changes, a filter resolution of 20 Å was used. Large changes in filament curvature and α-catenin occupancy were observed along the first variability component. Filament segments from the 0^th^ to the 25^th^ percentile, featuring high α-catenin occupancy, were selected and refined in RELION to produce an asymmetric reconstruction.

### Filament helical parameter measurements

Measurements were performed using our previously described approach^32^. Briefly, canonical F-actin protomers (PDB: 8D13) were rigid-body fitted into each reconstructed map and combined into a single model, and three copies of the model were stitched together to minimize edge artifacts. Only the central protomers from a single model were reported. A 3D spline is fit to the filament axis of each model. Rise is measured by computing the path length along the central axis between neighboring protomers.

Twist is determined using the Frenet-Serret frame of reference defined by the orthonormal basis of the unit tangent, unit normal, and unit binormal vectors along the length of the spline. The frame is then rotated along the normal-binormal plane. Twist is measured along the short pitch helix between consecutive protomers. The models analyzed in this work exhibited substantial lattice deformations that required minor adjustments to reliably measure the filaments’ helical parameters. The measurement approach was updated to account for a variable radius of the filaments. During fitting of the filament axis spline, the radius was allowed to vary along the filament length and was fit using a separate univariate spline; this tended to suppress spurious twist deviations. Additionally, the modest resolution of the reconstructed maps, particularly at the filament edges, limited our confidence in protomer fitting. Consequently, for the consensus superhelical F-actin, the -ATP control reconstructions, and the 3DVA sorted α-catenin-bound F-actin map, three independent manual rigid-body fittings were performed to provide confidence intervals for the helical parameter measurements.

### Quantifying actin subdomain rearrangements

First, 25 copies of a canonical protomer model (PDB: 8D13) were rigid-body fit into each map. These 25 protomers were combined into a single PDB for each map, and these combined models were subjected to Phenix geometry minimization with default parameters of 500 maximum iterations and 5 macro cycles in order to remove clashes. These minimized PDBs were then input to ISOLDE^45^ and hydrogen atoms were added. Secondary structure distance restraints were imposed on each actin subunit for the following residue ranges: 7-35, 35-72, 72-147, 340-377, 147-183, 272-340, 183-272, based on previously defined subdomain boundaries^46^. Torsional and distance restrains were then imposed on the entire secondary structure of each protomer. ISOLDE simulations were run with a weight of 30 x 1000 kJ mol^-1^ (map units)^-1^ Å^3^. The temperature was set to 100 K for 10 minutes of simulation, then slowly decreased to 0 K. Subdomain movement magnitudes and directions were determined as previously described^32^. Briefly, protomers were aligned to a reference protomer model (PDB: 8D13) and the displacement vectors between C_α_s of the flexibly fit protomers and the reference protomer were computed. Lastly, the average of the displacement vectors was calculated for each subdomain defined above.

### Molecular graphics and data analysis

Molecular graphics were prepared with ChimeraX^13^. Unless otherwise noted, plotting and statistical analysis was performed with GraphPad Prism. Python codes were prepared with the assistance of ChatGPT 4.0.

